# The extracellular matrix protein agrin is essential for epicardial epithelial-to-mesenchymal transition during heart development

**DOI:** 10.1101/2020.09.25.313742

**Authors:** Xin Sun, Sophia Malandraki-Miller, Tahnee Kennedy, Elad Bassat, Konstantinos Klaourakis, Jia Zhao, Elisabetta Gamen, Joaquim Miguel Vieira, Eldad Tzahor, Paul R. Riley

**Affiliations:** Burdon-Sanderson Cardiac Science Centre, Department of Physiology, Anatomy and Genetics, University of Oxford, Oxford OX1 3PT, UK; British Heart Foundation - Oxbridge Centre of Regenerative Medicine, CRM, University of Oxford, Oxford OX1 3PT, UK; Department of Molecular Cell Biology, Weizmann Institute of Science, Rehovot 76100, Israel; Research Institute of Molecular Pathology, Vienna Biocenter, Campus-Vienna-Biocenter 1, 1030 Vienna, Austria; Department of Cellular and Physiological Sciences, Life Science Institute, The University of British Columbia, 2350 Health Sciences Mall, RM.5320, Vancouver, BC, Canada, V6T 1Z3

**Keywords:** Epicardium, EMT, ECM, Agrin, Dystroglycan, Golgi apparatus, WT1

## Abstract

During embryonic heart development, epicardial cells residing within the outer layer of the heart undergo epithelial-mesenchymal transition (EMT) and migrate into the myocardium to support and stimulate organ growth and morphogenesis. Disruption of epicardial EMT results in aberrant heart formation and embryonic lethality. Despite being an essential process during development, the regulation of epicardial EMT is poorly understood. Here we report EMT on the epicardial surface of the embryonic heart at subcellular resolution using scanning electron microscopy (SEM). We identified high- and low-EMT regions within the mesothelial layer of the epicardium and an association with key components of the extracellular matrix (ECM). The ECM basement membrane-associated proteoglycan agrin was found to localize in the epicardium in regions actively undergoing EMT. Deletion of agrin resulted in impaired EMT and compromised development of the epicardium, accompanied by down-regulation of the epicardial EMT regulator WT1. Agrin enhanced EMT in human embryonic stem cell-derived epicardial-like cells by decreasing β-catenin and promoting pFAK localization at focal adhesions. In addition, agrin promoted the aggregation of its receptor dystroglycan to the Golgi apparatus in murine epicardial cells and loss of agrin resulted in dispersal of dystroglycan throughout the epicardial cells in embryos, disrupting basement membrane integrity and impairing EMT. Our results provide new insights into the role of the ECM in heart development, and implicate agrin as a critical regulator of EMT, functioning to ensure dystroglycan connects signals between the ECM and activated epicardial cells.

**Summary statement:** The basement membrane-associated proteoglycan agrin regulates epicardial epithelia-to-mesenchyme transition (EMT) through dystroglycan localizing on the Golgi apparatus. This ensures ECM and cytoskeletal connectivity and mechanical integrity of the transitioning epicardium and has important implications for the role of the extracellular matrix (ECM) in heart development.

## Introduction

The formation and growth of the mouse embryonic heart is a highly dynamic process that includes heart tube elongation and looping, chamber septation and growth. Early specified cardiac progenitors in the linear heart tube rapidly proliferate and differentiate, followed by the incorporation of distinct progenitors from the second heart field to support the structure and function of the mature 4-chambered heart (reviewed by Meilhac and Buckingham, 2018; Günthel et al., 2018). Other cell lineages such as the epicardium, endocardium and neural crest also contribute cells, including fibroblasts, endothelium and vascular smooth muscle to the developing heart (reviewed by Gise and Pu, 2012; Meilhac and Buckingham, 2018). In addition, proper deposition and modification of the extracellular matrix (ECM) is required for effective attachment, migration, and differentiation of the various cardiovascular cell types. However, the ECM has received comparatively less research-focus and, as a consequence the molecular determinants regulating the interactions between distinct cell compartments and matrix in the embryonic heart remain poorly understood.

ECM is the protein scaffold that not only supports cell attachment but also acts as a reservoir for signaling molecules (reviewed by Hynes, 2009, Hynes, 2014). The ECM dynamically interacts with cells to regulate their behavior and, in turn, cells feedback by modifying the matrix to adapt to their changing cell fate and condition the local environment. ECM mainly comprises of extensive networks of collagen and laminin, with basement membrane attachments mediated by proteoglycans and glycoproteins (reviewed by Rozario and DeSimone, 2009; Bonnans et al., 2014, Hynes, 2014, Walma and Yamada, 2020). The importance of ECM in various physiological processes has been the focus of a number of previous studies (reviewed by Larsen et al., 2006; Neill et al., 2015). During heart development, the ECM has been implicated in mesendoderm cell fate decisions, trabeculation and valve formation (Cheng et al., 2014; Monte-Nieto et al., 2018; Gunawan et al., 2019). Within the epicardium, the layer of mesothelial cells covering the outer surface of the myocardium which is essential for heart development, ECM components have been reported to contribute to tertiary structures within the mesothelium and the establishment of a stem cell-like niche (Balmer et al., 2014).However besides maintaining structural integrity, regulatory roles for the ECM within the epicardium and more generally during heart development have not been studied to-date.

In mouse, the epicardium is derived from the proepicardium (PE), which is a transient embryonic structure attached at the base of venous inflow tract. At embryonic day (E) 9.5 PE cells migrate to cover the surface of the myocardium forming an outer cell layer by E11.5. A subset of epicardial cells undergo epithelial-to-mesenchymal transition (EMT) and invade into the underlying myocardium, with epicardium-derived cells (EPDCs) then differentiating into multiple cardiovascular cell types including fibroblasts and vascular smooth muscle cells to support coronary vessel development and growth of the embryonic heart (reviewed by Riley 2012, Simoes and Riley, 2018, Cao and Poss 2018). Disruption of epicardial development leads to mid-gestation lethality, with mutant mice exhibiting underdeveloped hearts (Yang et al., 1995; Kwee et al., 1995). For instance, loss of the master epicardial cell regulator, the transcription factor Wilms’ tumor 1 (WT1) resulted in defective epicardial EMT, hypoplastic myocardium and embryonic death (Moore et al., 1999; Zhou et al., 2008; Von Gise et al., 2011). WT1 along with several other factors, including fibroblast growth factor (FGF), transforming growth factor beta (TGFβ), and thymosin beta 4 (Tβ4), are also critical for the formation of the epicardium and maintenance of epicardial integrity, suggesting possible roles for the ECM of the basement membrane of the epicardium (Vega-Hernandez et al., 2011; Compton et al, 2007; Zhou et al., 2012; Smart et al., 2007). The precise spatial-temporal events supporting epicardial EMT have not been described in detail and how the ECM and associated intracellular signals interact and enable the transition from an intact epithelium and epicardial cell migration is unknown.

Agrin is an important component of the ECM with broad signal-mediating potential to ensure connectivity between cells and the basement membrane. It is a heparan-sulphate proteoglycan (HSPG) broadly expressed in muscle, neuron, blood and endothelial cells (Stone et al., 1995). Agrin was first identified as a protein that can promote the aggregation of acetylcholine receptors on the muscle-neuron surface through muscle-synapse kinase (MuSK) (Nitkin et al., 1987; Gautam et al., 1996; Lin et al, 2001). More recently, it has been implicated in cancer progression and invasion (Charkroboty et al., 2015; Rivera et al., 2018). In the brain endothelium, agrin deposits in the basement membrane, ensheathing blood vessels and contributing to blood-brain barrier (BBB) maturation (Steiner et al., 2014). Mechanistically, agrin transduces tissue rigidity signals and stabilizes the Yes-associated protein (YAP) (Chakraborty et al., 2017). Recently it has been implicated as an important contributor to cardiomyocyte stiffness and turn-over in the context of heart regeneration and repair in both mice and pigs (Basset et al., 2017; Baehr et al., 2020). However, the role of agrin in development per se and more specifically within the embryonic heart has not previously been described.

Here, we reveal for the first time a requirement for agrin in the developing heart as an integral link between the ECM and epicardial development. We employed high-resolution imaging using combined scanning electron microscopy (SEM) and confocal microscopy to demonstrate that the embryonic epicardium comprises a highly heterogenous morphology with evidence of epicardial cells undergoing regionalized EMT. ECM components, laminin and integrin α4, agrin and its receptor dystroglycan were associated with active regions of EMT in the developing epicardium. Loss of agrin resulted in pleiotropic defects, including compromised epicardial cell proliferation, abnormal deposition of epicardial ECM and decreased EMT. Mechanistically we reveal that exogenous agrin promoted EMT and activated the integrin-focal adhesion kinase (FAK) signalling pathway, in a model of human embryonic stem cell-derived epicardial cells, and that aggregation of dystroglycan to the Golgi apparatus acts as the signal transduction pathway for agrin in epicardial cells, and is required for maintaining basement membrane and cytoskeletal connectivity during EMT. Taken together, our study identifies the ECM component agrin as a a critical regulator of epicardial EMT and novel determinant of normal heart development.

## RESULTS

### Epicardial cells with distinct morphology and EMT status are present on the surface of the embryonic heart

To investigate changes in cell morphology during epicardial EMT, we initially examined the outer surface of the developing mouse heart at E13.5 and E14.5 by SEM (**Figure 1A-1H**). At E13.5, the ventral surface of the heart was found to be irregular (**Figure 1A; n=5**), with clusters of rosette-like structures separated by grooves being observed at higher-magnification (**Figures 1B-1D**). By comparison, the dorsal epicardial surface appeared to be smoother (**Figure S1A**), while two distinct cell morphologies could be observed on the epicardial surface: large flat-shaped cells connecting with neighboring cells with well-defined cell-cell borders (**Figure S1B, white arrow**) and loosely connected small pillar-shaped cells (**Figures S1B, S1C, white arrowhead**). The latter morphology-type was represented throughout the heart surface, with small clusters (**Figure 1D, white arrow**) being observed among patches of tightly-connected, large flat cells (**Figure 1D, white arrow head**). These two cell morphologies were also observed on the epicardial surface at E14.5 (**Figure 1E**-**H; n=6**). The large flat surface cells exhibited finger-shaped protrusions mainly along the cell-cell borders (**Figure 1G, white arrow; Figure S1B, white arrow**), whereas such protrusions were distributed on the apical and lateral sides of small pillar-shaped surface cells (**Figure 1H, white arrow; Figure S1C**). Importantly, the changes in cell morphology observed are consistent with the definition of cells undergoing EMT, i.e. transition from a large flat shape into a small pillar-like structure (Yang et al., 2020).

**Figure 1.**
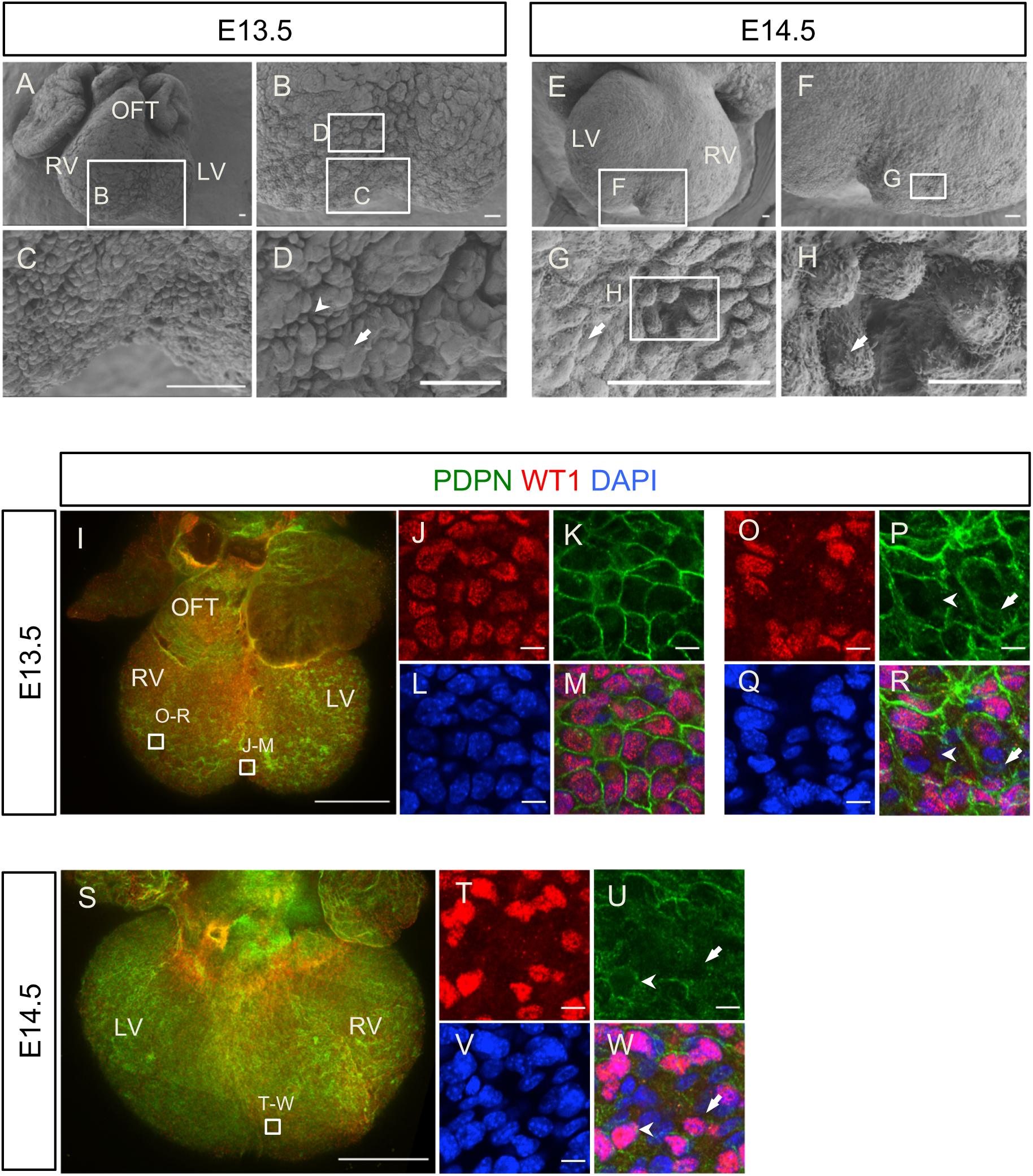
The developing epicardium has active regions of epithelial-mesenchymal transition (EMT) and exhibits morphological heterogeneity. (A-H) The surfaces of E13.5 and E14.5 hearts under Scanning electron microscopy (SEM). (A) The ventral aspect of an E13.5 heart. (B) Magnified view from the inset box in (A) showing the surface covering the ventricular wall and the interventricular septum (IVS). (C, D): Magnified view of inset boxes in (B). (E) the dorsal aspect of an E14.5 heart. (F) Magnified view of the inset box in (E) showing the surface of the ventricle wall and IVS. (G) Magnified view of the inset box in (F). (H) Magnified view of the inset box in (G). Note the distinct cell morphologies including flat and tightly connected cells (white arrow in G) and cells detached from each other suggesting undergoing EMT (white arrow in H). (I-R) Whole-mount immunofluorescent staining for epicardium markers WT1 (red), podoplanin (PDPN, green) and DAPI (blue) in an E13.5 heart shown at ventral aspect. (J-M) magnified view of the inset box in (I) at the ventricular septation groove. (O-R) magnified view of the inset box in (I) on the ventricle wall. Note the various shapes of WT1 positive nuclei and podoplanin labeled cell boundaries. (S-W) Whole-mount immunofluorescent staining for WT1 (red), podoplanin (green) and DAPI (blue) in an E14.5 heart at dorsal aspect. (T-W) magnified view of the inset box in (S) near the ventricle septum groove. Scale bar: A-G: 50 μm. H: 10 μm. I and S: 500 μm. J-R, T-W: 10 μm. LV: left ventricle. RV: right ventricle. OFT: outflow tract.

To further characterize the epicardial surface during heart development, we performed whole-mount immunofluorescence staining using antibodies against the well-established epicardial markers Podoplanin (PDPN) and WT1 (Mahtab et al., 2008; Moore et al., 1999; Zhou et al., 2008). PDPN immunoreactivity was detected on the lateral borders of epicardial cells, while WT1 expression was restricted to their nuclei (**Figures 1I-1W**). We observed large, round WT1^+^ nuclei with podoplanin-positive cell borders near the ventricular septum groove (**Figures 1J-M**) and epicardial cells undergoing EMT within the epithelium in this region (**Figures 1F, 1G**). On the ventricular wall, the WT1^+^ nuclei varied in shape as round, oval or rod-like. Some of the cells in the epicardium had down-regulated WT1 or lacked WT1 in their nuclei, although PDPN was detected in their membranes (**Figures 1O, 1P, 1R**, white arrow), suggesting they were differentiating epicardial cells. Interestingly, we observed small regions with multiple WT1^+^ and WT1^-^ nuclei, but no PDPN^+^ cell borders separating them (**Figures 1P, 1R**, white arrowhead). We hypothesized that absence of podoplanin in these regions may be associated with EMT regions where cell-cell connections are lost, as observed in our SEM analyses (**Figures S1B, S1C**). At E14.5, we observed similar epicardial cell morphologies (**Figures 1S-1W**). WT1^+^ epicardial cells enclosed with defined PDPN^+^ borders were evident (**Figures 1U, 1W**, white arrowhead). alongside regions in which PDPN was low and discontinuous with both WT1+ and WT1-nuclei (**Figure 1U, 1W**, white arrow) and, as for E13.5, potentially labelling active regions of epicardial EMT.

On the outflow tract, SEM revealed that epicardial cells were small and round and tightly packed together (**Figure S2A)**. Whole-mount immunofluorescent staining confirmed the cells at this location were WT1 and PDPN positive (**Figures S2B-D**). In contrast, cells at the junction of the outflow tract with the chambers were large and flat (**Figure S2A)**. These cell shape differences were also observed by immunofluorescent staining for PDPN (**Figure S2C**). In summary, combining SEM and immunofluorescent staining, we observed regional and heterogeneous cell morphology within the developing epicardium, not previously described, and we attribute this to representing the EMT status of the constituent epicardial cells.

### Active epicardial EMT regions have a distinct extracellular matrix (ECM) composition

Since EMT is associated with dynamic basement membrane breakdown and establishment, we investigated the distribution of key ECM components: integrins α4 and β1, laminin and fibronectin in the developing epicardium, using whole-mount immunofluorescent staining. Integrin α4 is an epicardial-specific integrin subunit (Yang et al., 1995; Kirschner et al., 2006) and integrin β1 is the beta-subunit predominantly expressed in the developing heart (Ieda et al., 2009). Laminin is a major component of the basal membrane (reviewed by Yurchenco, 2015). Fibronectin is a high-molecular weight glycoprotein and serves as a receptor for various integrins (reviewed by Pankov and Yamada, 2002) and is secreted by epicardial cells and necessary for heart regeneration in zebrafish (Wang et al., 2013). These ECM proteins were all detected on the surface of the embryonic heart at E14.5 (**Figures S3A-3D**), with closer examination showing enrichment in regions proximal to the forming coronary vasculature (**Figures S3E-S3H**).

To investigate further the association of ECM components with epicardial cell delamination and migration into the subepicardial space and underlying myocardium, we analyzed serial sagittal sections of hearts at E13.5 (**Figure 2**). We focused on a region in the ventral side with clusters of WT1^+^ cells migrating from the epicardium to the myocardium (**Figures 2A-2F**). In this region, WT1^+^ cells were tightly packed together into multiple layers. Moreover, clusters of WT1^+^ cells with large and round nuclei were found in the underlying myocardium (**Figures 2A, 2C** white arrows, **2D, 2F**). Here, integrin α4 expression was found to be weak and discontinuous around migrating WT1^+^ cells (**Figures 2A, 2B, 2D, 2E**). By contrast, the integrin α4 signal was clearly visible and continuous on the epicardium in other regions of the same sections, where WT1^+^ cells were mainly localized within the epicardium (**Figures 2A, 2B, 2C**, white arrowhead). Comparatively fewer WT1^+^ cells were found migrating into the myocardium in these integrin α4-enriched epicardial regions (**Figures 2A-2C**). This observation suggests that down-regulation of integrin α4 is associated with active EMT.

**Figure 2.**
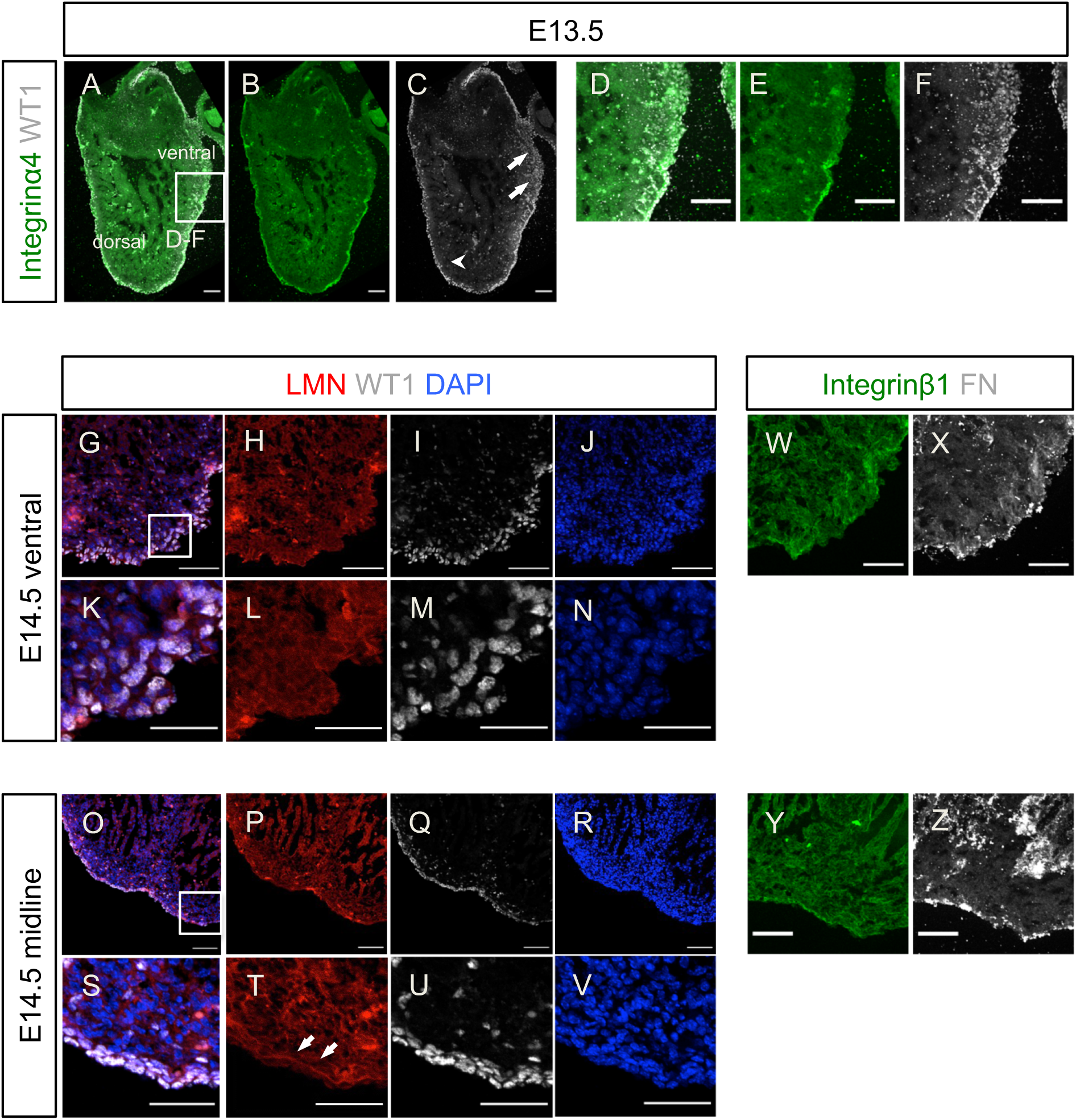
Active EMT regions within the embryonic epicardium are associated with extracellular matrix (ECM) components. (A-C) Immunofluorescent staining of a sagittal section of an E13.5 heart for WT1 (white), Integrin α4 (green) and DAPI (blue) showing clusters of WT1^+^ cells migrating into the myocardium and down-regulated Integrin α4 around these cells. (D-F) magnified view from the inset box in (A). (G-V) Immunofluorescent staining for laminin (red), WT1 (white) and DAPI (blue) on serial sections from one E14.5 heart. (G-N) a section near the ventral surface. (O-V) a section near the dorsal-ventral midline. (K-N) is magnified view of the inset box in (G). (S-V) is magnified view of the inset box in (O). Laminin in the region shown in (H and L) is discontinuous. In this region, clusters of WT1^+^ cells actively undergoing EMT is shown in (I and M). In the other section, The laminin forms a continuous expression domain along the epicardium (arrows in T). WT1^+^ cells are tightly associated with the epicardium (Q and U). W and X: immunofluorescent staining of a neighbor section of (G) for Integrinβ1 (green) and fibronectin (FN, white). Y and Z: immunofluorescent staining of a neighbor section of (O) for Integrinβ1 (green) and fibronectin (white). Neither integrinβ1 nor fibronectin showed association with EMT of WT1^+^ cells. Scale bar: A-F: 100 μm. G-Z: 50 μm.

By examining a series of long-axis serial sections from hearts at E14.5, we observed active EMT regions (**Figures 2G, 2K**) to be proximal to the ventral surface, whereas low EMT regions (**Figures 2O, 2S**) tended to be distal, closer to the heart midline. Importantly, in the active EMT regions, deposition of laminin around WT1^+^ cells was low and uneven (**Figures 2K, 2L**). In this region, the majority of WT1^+^ nuclei were large and round in shape (**Figures 2M**). By contrast, in low EMT regions, laminin formed a continuous sheet on the side of epicardial cells contacting the myocardium (**Figure 2O, 2P, 2S, 2T**, white arrows). WT1^+^ nuclei were rod-shaped and tightly attached within the laminin-marked basal membrane (**Figure 2O, 2Q, 2S, 2U** and fewer WT1^+^ cells were detected in the adjacent myocardium (**Figure 2Q, 2U**). Integrin β1 expression was detected in the epicardium and myocardium (**Figures 2W, 2Y**), with no obvious difference in the high (**Figure 2W**) versus low (**Figure 2Y**) epicardial EMT regions. Likewise, the punctate expression pattern of epicardial fibronectin showed no difference in high versus low-EMT regions (**Figure 2X, 2Z**), suggesting that the distribution of integrin β1 and fibronectin is not associated with epicardial EMT.

### Agrin is expressed in the epicardium and regulates epicardial EMT in the developing heart

To further characterize the ECM components associated to epicardial EMT, we next focused on agrin, an important component of the basement membrane that can be bound to the cell membrane or secreted (reviewed by Bezakova and Ruegg, 2003). Using whole-mount immunofluorescent staining, we investigated the expression of agrin in the developing epicardium from E13.5 to E16.5, focusing on regions proximal to the interventricular septum (**Figures 3A, 3H, 3O**), which appeared enriched for cells undergoing EMT as detected by SEM (**Figure 1**). At E13.5, agrin was detected in the WT1^+^ epicardial layer in a sparse, punctate pattern along the cell borders (**Figures 3B-3F**, white arrows), colocalizing with PDPN (**Figures 3C, 3F**). At E14.5, among the tightly packed WT1^+^ PDPN^+^ cells near the septum (**Figures 3I, 3J, 3K**), agrin-expressing puncta were more dense (**Figures 3J, 3M**). By E16.5, many epicardial cells had prominently down-regulated or lost WT1 expression coincident with completion of EMT and subsequent differentiation (**Figures 3P, 3Q, 3S**). Likewise, the punctate expression pattern of agrin in epicardial cells was also reduced (**Figures 3Q, 3T**, white arrows), suggesting agrin is associated with epicardial EMT.

**Figure 3.**
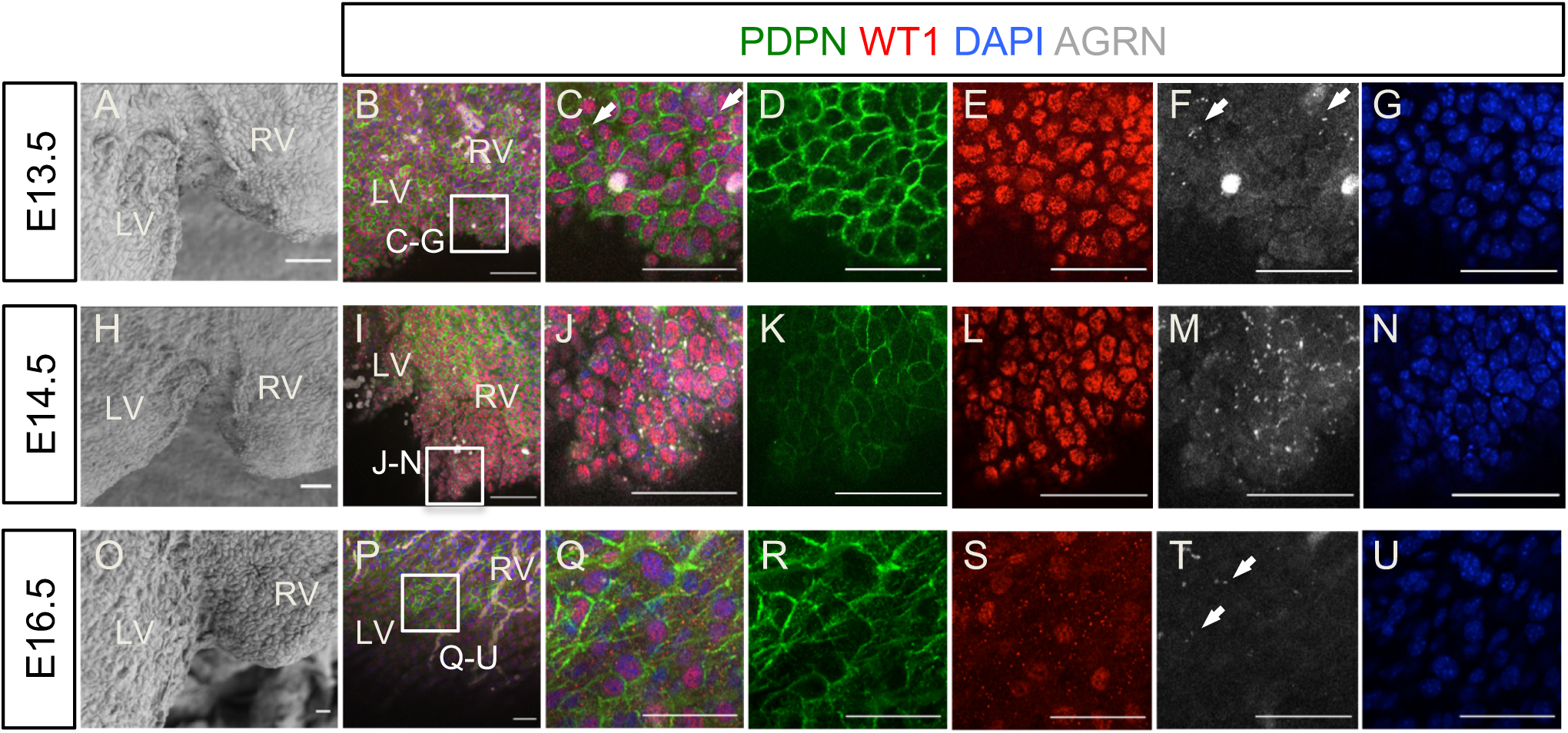
Localization of agrin in the developing epicardium. (A, H, O): SEM showing the area near the interventricle septum of E13.5, E14.5 and E16.5 hearts, respectively. (B-G, I-N, P-U): Whole-mount immunofluorescent staining of an E13.5, E14.5 and E16.5 heart for podoplanin (PDPN, green), WT1 (red), agrin (white) and DAPI (blue). (C, J, Q) are magnified views from the inset boxes of (B, I, P). Agrin localizes among epicardial cells in close proximity of podoplanin (C, F, white arrows). The punctate localization of agrin increased at E14.5 (J and M) and decreased at E16.5 (Q, T). Scale bar: 50 μm. LV: left ventricle. RV: right ventricle.

Given that a requirement for agrin during heart development has not been previously described, we generated global agrin knockout mutants by crossing *Agrn*^flox/flox^ (Harvey S et al, 2007) with *Pgk-Cre* mice (Lalleman Y., et atl., 1998). The resulting *Agrn*^+/-^ progenies were intercrossed to generate *Agrn*^-/-^ embryos, henceforth referred to as agrin KO. SEM was used to initially characterize the epicardial surface of agrin KOs versus control littermate hearts at E14.5 and E16.5. Whilst wild-type epicardial cells displayed multiple protrusions along the cell-cell border and on the cell surface, the density of protrusions appeared reduced on the epicardium of agrin KOs (**Figures 4A, 4B**, white arrows). This phenotype was also observable in hearts at E16.5 (**Figures 4C, 4D**, white arrows). Next, we assessed proliferation levels in agrin KO hearts by immunostaining against the phospho-histone H3 marker (PH3). At E14.5, agrin KO hearts presented an overall decrease in PH3^+^ cells (**Figures 4E-H**), with the number of PH3^+^ epicardial cells being significantly lower in KO hearts compared to littermate controls (30.4% ± 3.95% PH3^+^ epicardial cells in wild-type versus 10.3% ± 0.42% PH3+ epicardial cells in agrin KO; n=4, p=0.00095; **Figure 4I**). Moreover, fluorescent immunostaining against WT1 revealed that there were significantly fewer WT1^+^ cells on the epicardium as well as in the myocardium (epicardium: 52.9% ± 8.27% WT1^+^ cells in wild-type versus 32.5% ± 7.47% WT1^+^ cells in agrin KO, n=3, p=0.018; myocardium: 1.53 ± 0.55 WT1^+^ cells per artificial unit (1×10^4^ DAPI^+^ pixels) in wild-type versus 0.55 ± 0.24 WT1^+^ cells per artificial unit in agrin KO, n=3, p=0.02; **Figures 4J-4O**), suggesting that formation of the epicardium and epicardial cell migration are agrin-dependent. To further validate these results, we investigated PDPN immunoreactivity in agrin KO versus control littermates, and found that expression levels were low and discontinuous in the epicardium of mutant hearts at E14.5 (compare **Figures 4P and 4Q**). Likewise, at E16.5 whole-mount immunofluorescent staining revealed that podoplanin expression was down-regulated on the epicardial surface of agrin KO hearts (**Figures 4R-4U**), with the epicardial cell-cell border barely discernible (compare **Figures 4T with 4U**). Furthermore, the growth and patterning of PDPN-expressing lymphatic vessels in the subepicardial layer appeared defective in KO versus wild-type hearts (compare **Figures 4R with 4S**). The expression of WT1 was also decreased in the outer surface of KO hearts at E16.5 (**Figures 4V, 4W;** note that these panels are from the same region as in **Figures 4T and 4U**). At this stage, 18.17% ± 3.21% of cells in the epicardium of controls expressed WT1^+^, while in the agrin KO only 5.29% ± 1.36% of cells were WT1^+^ (n=3, p=0.02; Figure **4X**). In summary, loss of agrin resulted in abnormal epicardial development and EMT.

**Figure 4.**
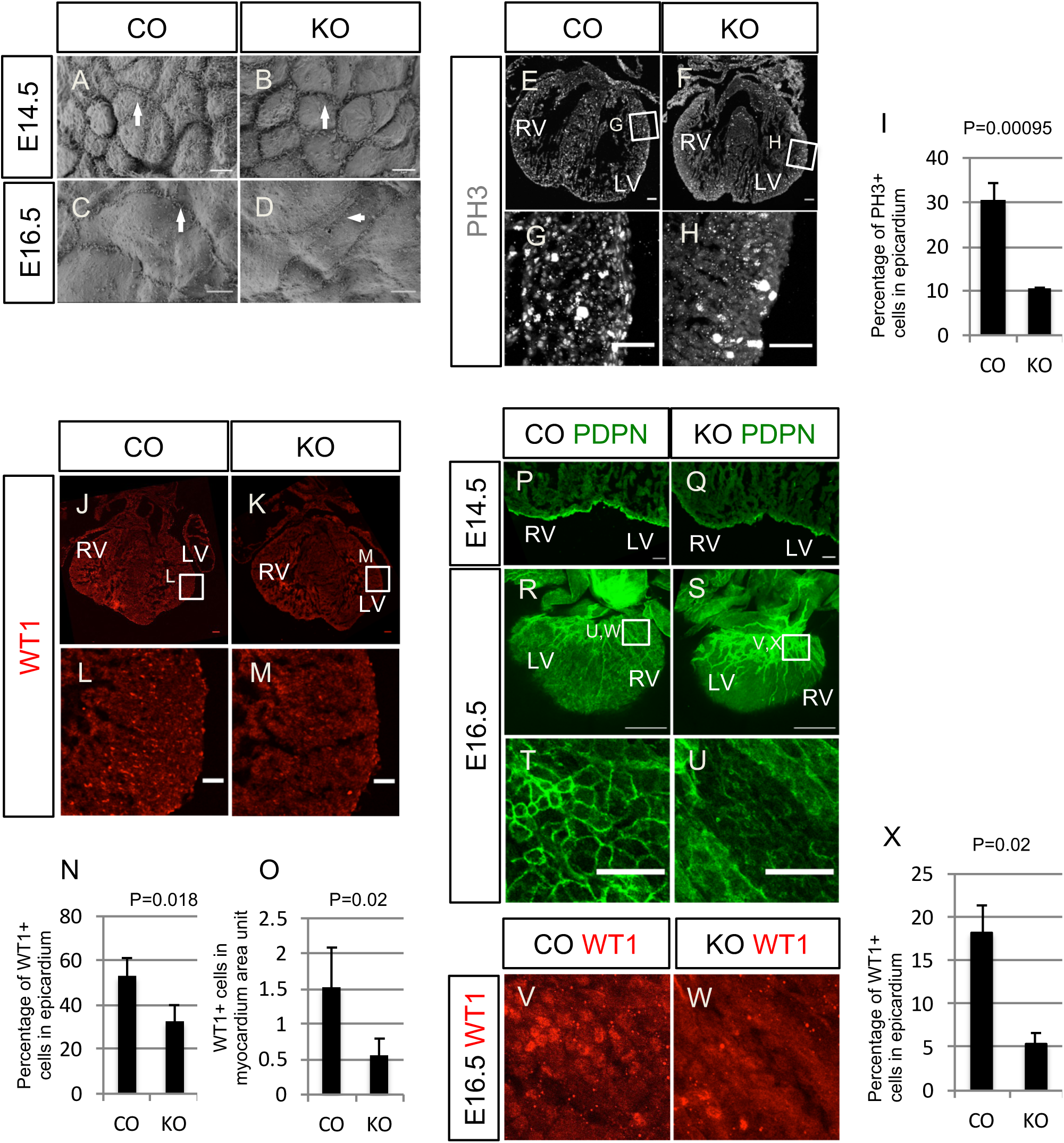
Loss of agrin results in developmental defects specifically in the epicardium. (A-D) Epicardial cells on the surface of E14.5 (A) and E16.5 (C) littermate controls and agrin KO hearts (B and D) hearts under SEM. (E-H) immunofluorescent staining for the proliferation marker phosphor-histone H3 (PH3; white) in sections of E14.5 hearts of control (C) and agrin KO (D) hearts. (G, H) are magnified views from the inset boxes in C and D. (I) Quantification of PH3^+^ epicardial cells in controls and agrin KO hearts (percentage of total DAPI in the epicardium). Data represent mean ± SEM. N=4 hearts per group. Significant differences (p value) is calculated using an unpaired, two-tailed Student’s *t*-test. (J and K) Immunofluorescent staining for WT1 (red) in E14.5 heart sections of the littermate control and agrin KO. (L and M) are magnified views from inset boxes in (J and K) showing WT1^+^ cells on the epicardium and in the myocardium. (N) Quantification of WT1^+^ cells in the epicardium of the controls and KO (percentage of WT1^+^ nuclei in total DAPI in epicardium). (O) Quantification of WT1^+^ cells in myocardium (percentage of WT1^+^ nuclei in total myocardial DAPI area). Data represent mean ± SEM. N=4 hearts per group. Significant differences (p-value) is calculated using an unpaired, two-tailed Student’s *t*-test. (P, Q) Immunofluorescent staining for podoplanin (PDPN, green) in E14.5 control (P) and agrin KO (Q) sections. (R, S) Whole-mount immunofluorescent staining of E16.5 control (R) and agrin KO (S) hearts for podoplanin (green) labeling cardiac lymphatic vessels and epicardial cells. (T and U) are magnified views of the inset boxes in (R and S). Note the down-regulated epicardial podoplanin in agrin KO hearts. (V, W) Whole-mount immunofluorescent staining for WT1 (red) of the same hearts in (R, S) at the same area of (T, U). (X) Quantification of percentage of WT1^+^ cells in epicardium of E16.5 controls and agrin KO (percentage of WT1^+^ nuclei compared to total DAPI in epicardium. Data represent mean±SEM. N=3 hearts per group). Significant differences (p-value) was calculated using an unpaired, two-tailed Student’s *t*-test. Scale bar: LV: A-D: 5 μm. E-H, J-M: 50μm: G, H, L, M: 50μm. Q, R: 10 μm. S, T: 500μm. U, V, W, X: 50μm. LV: left ventricle. RV: right ventricle.

Since the disruption of epicardial development may affect the formation of coronary vasculature (Gise et al, 2011), we investigated whether loss of agrin also resulted in vascular defects. At E14.5, agrin KOs exhibited shortening of the superficial coronary veins as depicted by endomucin (EMCN) staining in the dorsal aspect of the heart (**Figures S4A, S4B**, white arrows). This defect was also observed in hearts at E16.5 (**Figures S4C, S4D**, white arrows) and by E18.5, the coronary vessels in agrin KO hearts appeared to have lost vessel integrity with PECAM^+^ endothelial cells failing to form a fully enclosed lumen (**Figures S4E-S4H**, white arrow).

Next we characterized the expression of other ECM proteins in the agrin KO hearts. We found that epicardial integrin α4 was down-regulated in agrin KO hearts (**Figures 5A, 5B**) whereas, in contrast, integrin β1 was unchanged in the mutants (**Figures 5C, 5D**). Collagen I and Laminin, both important components of basal membrane ECM, were expressed at lower levels in agrin KO hearts at E14.5, in particular within the myocardial compartment (**Figures 5E-5H**). These results suggest that agrin is important for deposition, assembly or turnover of ECM components in relation to epicardial cell basement membrane connections and ongoing EMT. Of significance, we found cytosolic β-catenin was upregulated in mutant epicardial cells and in cells immediately adjacent to the epicardium, suggesting these cells have tighter connections than those in hearts from littermate controls compromising their EMT potential (**Figures 5I, 5J**). We measured and compared the average intensity of collagen I and laminin in embryonic heart sections of E14.5 littermate controls and agrin KO which revealed both were down significantly regulated in agrin KO hearts (collagen I: 135.23 ± 8.43 in controls vs 106.54 ± 7.38 in agrin KO, n=3, p=0.011; laminin: 136.57 ± 10.2 in controls vs 113.29 ± 9.59 in agrin KO, n=3, p=0.0009. Figure **5K, 5L**).

**Figure 5.**
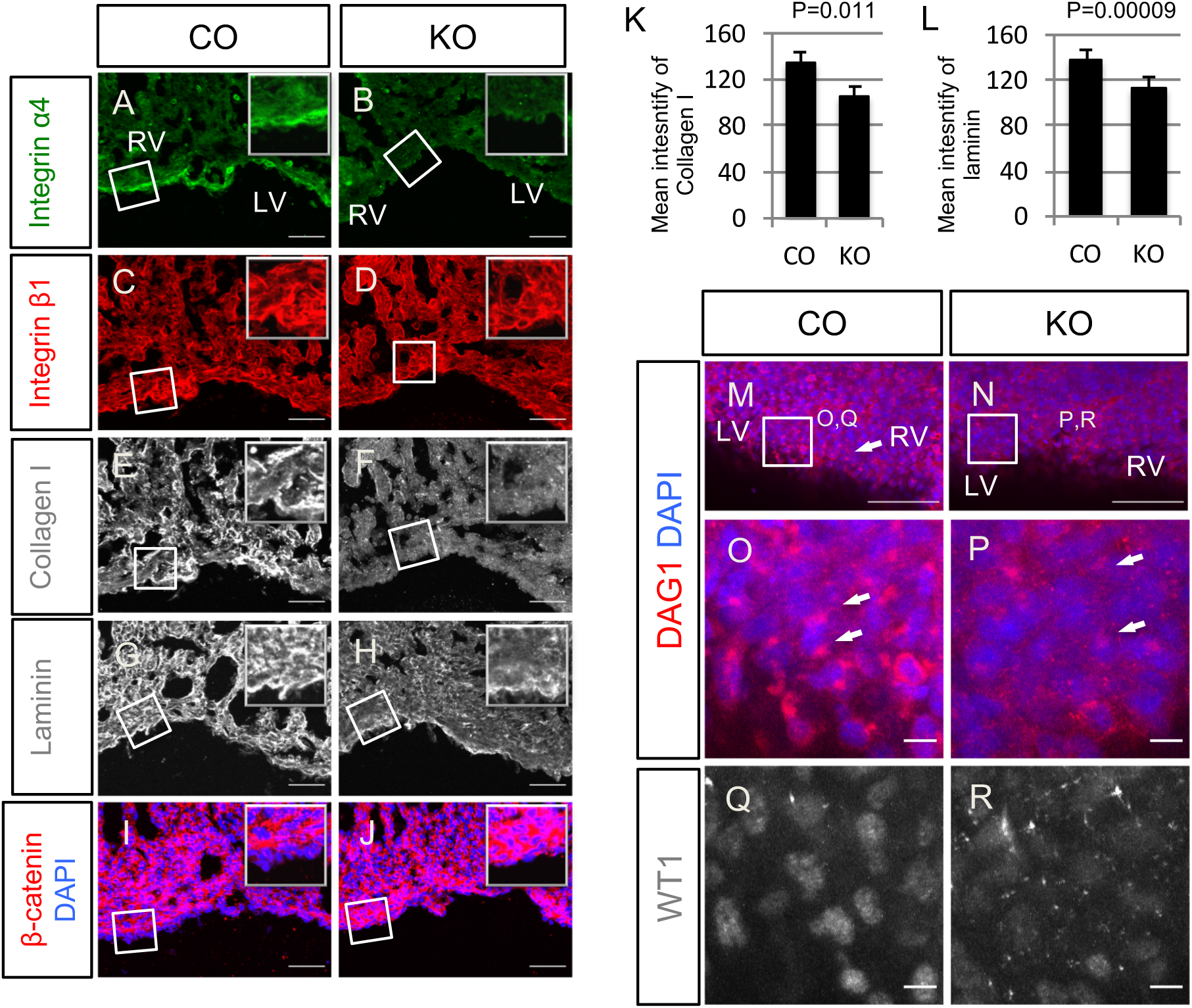
Deletion of agrin compromises ECM deposition and dystroglycan aggregation in embryonic hearts. Immunofluorescent staining of Integrin α4 (A-B), Integrin β1 (C-D), Collagen I (E-F), Laminin (G-H) and β-catenin (I-J) in serial sections of embryonic hearts of E14.5 littermate control (A, C, E, G, I) and agrin KO (B, D, F, H, J). Note C, E are from the same region of a control section, D,F are from the same region of a agrin KO section. (K) Quantification of mean intensity of collagen I in embryonic heart sections of E14.5 controls and agrin KO. (L) Quantification of mean intensity of laminin in embryonic heart sections of E14.5 controls and agrin KO. Data represent mean ± SEM. N=3 hearts per group. Significant differences (p-value) was calculated using an unpaired, two-tailed Student’s *t*-test. (M-P) Wholemount Immunofluorescent staining using DAG1 antibody to show localization of dystroglycan in the epicardium of a littermate control (M, O) and agrin KO heart (N, P). (O and P) are magnified view of the inset boxes of (M and N), respectively, showing the dystroglycan puncta localizing in proximity with nuclei. (Q-R) Immunofluorescent staining of WT1 of the same regions in (O and P) showing down-regulation of WT1 in E16.5 agrin KO (R) compared with the littermate control (Q). Scale bar: A-J, M-N: 50μm. O-R: 10μm. LV: left ventricle. RV: right ventricle.

Dystroglycan is a glycoprotein connecting the extracellular matrix with intracellular actin (Ervasti and Campbell, 1993) and a known binding partner of agrin (Gee et al., 1994). Therefore, we investigated the distribution of dystroglycan in the embryonic heart using immunofluorescent staining with an antibody against DAG1. At E14.5, the DAG1 antibody detected dense puncta in close proximity to the nuclei of epicardial cells (**Figures 5M, 5O**, white arrows). By contrast, in agrin KO hearts dystroglycan-enriched puncta were either very weak or dispersed (**Figures 5N, 5P**, white arrows) and this in-turn correlated with low WT1 expression indicative of reduced epicardial activation and EMT (**Figures 5Q, 5R**).

To further characterize the impact of agrin-deficiency on EMT, we employed an epicardial explant model using E11.5 mouse hearts (Trembley et al., 2016). Epicardium-derived cells arising from agrin KO explants exhibited significantly reduced WT1 expression compared to wild-type explants (control 85% ± 6.2% vs KO 76% ± 4.5%, p=0.03, n=4; **Figures S5A-S5E**). Moreover, explants derived from agrin KO hearts exhibited less phalloidin-labeled stress fibers in the periphery of the outgrowth, supporting the hypothesis that agrin is required for epicardial EMT (**Figures S5C, S5D**). We next investigated the expression pattern of PDPN, as this glycoprotein is normally down-regulated in cells at the leading edge of epicardial sheets (mesenchymal-like cells), whilst remaining at high levels in the epithelial-like cells closer to the explant (Cao et al., 2017). Whilst PDPN levels were comparable between KO and control explants, its distribution within the cells differed; uniformly localized throughout the cell membrane and cytoplasm in controls, whilst enriched exclusively in a peri-nuclear pattern in mutant explant cells (**Figures S5F-S5I**, white arrows). Smooth muscle myosin heavy chain (SM-MHC) is a protein expressed in vascular smooth muscle cells thus used as an indicator for epicardial cell differentiation towards a smooth muscle fate. The incidence of SM-MHC expression in epicardial cells migrating from explants did not differ between agrin KO and control groups (**Figures S5J-S5L**), indicating that smooth muscle cell differentiation was not affected. Also, in the epicardial cells migrating from explants, there was no difference in the signal or localization of laminin between the agrin KO epicardial cells and the control cells (**Figures S5M, S5N**).

In summary, loss of agrin from explants resulted in down-regulation of epicardial markers (e.g. WT1, PDPN), defected ECM organization and compromised epicardial cell migration, but not their differentiation potential towards smooth muscle.

### Agrin promotes EMT in cultured epicardial cells

Our data revealed that loss of agrin impacts on mouse epicardial development, coronary vasculature formation and epicardial EMT. To further characterize the role of agrin in EMT and assess a potential conservation of this function in the human epicardium, we adopted an *in vitro* differentiation protocol generating epicardium-like cells from human embryonic stem cells (hESCs) (Iyer et al., 2015; **Figure 6A**). Following treatment with 5ng/ml TGFβ, a known inducer of EMT for 72hrs, the human epicardium-like cells presented clear morphological changes, such as enhanced stress fiber density (compare **Figures 6B** and **6C**). Importantly, we observed a similar phenotype after treating the cells with increasing concentrations of recombinant agrin (10ng/ml, 50ng/ml, 200ng/ml) (**Figures 6D-6F**). In addition to stress fiber density, we assessed the levels of phosphorylation of focal adhesion kinase (FAK) at tyrosine 397 (pFAK Y397), which connects the extracellular matrix signals and intracellular signaling through integrins and is a key regulator of cell migration, an important process in cells undergoing EMT (reviewed by Mitra et al., 2005, Larsen et al., 2006). Focal adhesion kinase (FAK) is a tyrosine kinase connecting extracellular matrix signals and intracellular signaling through integrin. It is a key regulator for cell migration. Phosphorylation of FAK at Tyrosine 397 (pFAK Y397) results its recruitment to focal adhesion and serving as signaling axis (reviewed by Mitra et al., 2005, Larsen et al., 2006). Treatment with 5ng/ml TGFβ enhanced the localization of pFAK Y397 at focal adhesions (**Figure 6H**). Similarly, agrin also induced focal adhesion localization of pFAK Y397, suggesting agrin had a similar effect as TGFβ in promoting human epicardium-like cells towards a mesenchymal fate (**Figures 6I-6K**).

**Figure 6.**
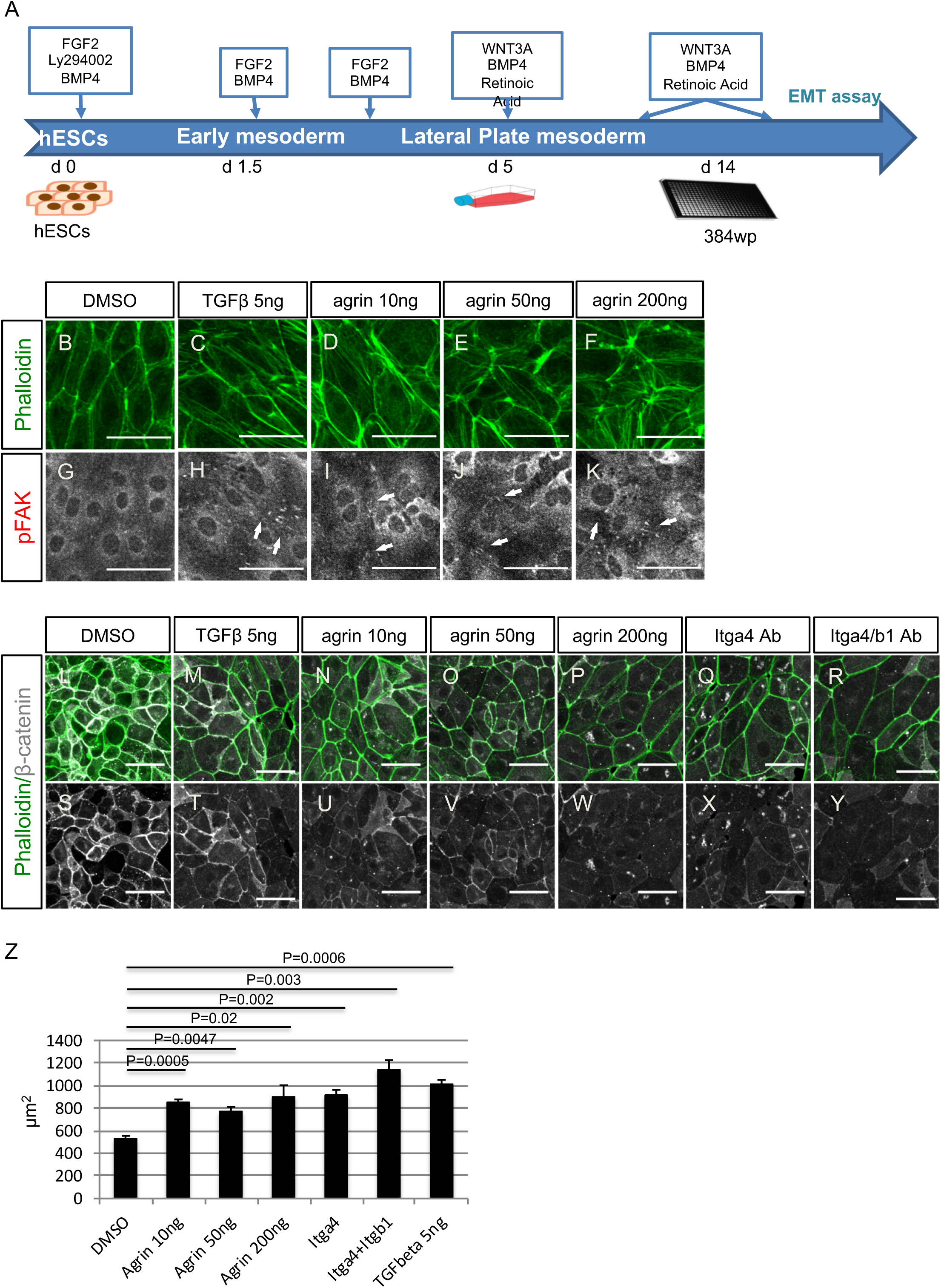
Agrin promotes EMT through enhancing pFAK and decreasing β-catenin. (A) A scheme showing the differentiation protocol of human epicardial cells (hEPDCs) from hESC. (B-F) Treatment of human epicardial-like cells with TGFβ and agrin resulted in enhancement of stress fibers (phalloidin, green). (G-K) immunofluorescent staining of cells in the same regions with (B-F) for pFAK (white). (L-Y) immunofluorescent staining for stress fibers (phalloidin, green) and β-catenin (white) on human epicardial-like cells treated with agrin or blocking antibody against integrin α4 and integring β1. Agrin induced a decrease in β-catenin expression at the cell-cell borders (N-P, U-W). Blocking antibody against epicardial integrin α4 and integrin β1 also decreased β-catenin (Q-R, X-Y). (Z) Addition of agrin enhanced cell size, indicating EMT. Data represent mean ± SEM. N=3 treatments per group. Significant differences (p value) was calculated using one-way ANOVA followed up by the two-tail student *t*-test between the two groups. All scale bars: 50μm.

In the *in vitro* model, the epithelial-like cells exhibited high numbers of cell-cell junctions (**Figures 6L, 6S**). Treatment with 5ng/ml TGFβ down-regulated the localization of β-catenin at the cell-cell junctions (**Figures 6M, 6T**); an equivalent down-regulation was also observed following addition of agrin (10ng/ml, 50ng/ml, 200ng/ml) consistent with the EMT promoting effect of agrin. The effect of agrin was dose-dependent and stronger than TGFβ at high dosage (**Figures 6N-6P, 6U-6W**). Incubation of the human epicardium-like cells with the blocking antibody Natalizumab (“Itga4 ab”), specific for the epicardially-expressed α4 integrin, also reduced β-catenin localization in cell-cell junctions (**Figures 6Q, 6X**). Combination of treatment with blocking antibodies against the epicardial specific α4β1 integrin had a stronger effect in decreasing β-catenin at cell-cell junctions (**Figures 6R, 6Y**). Addition of agrin or integrin blocking antibodies also significantly enhanced the cell size, phenocopying TGFβ treatment (**Figure 6Z**), consistent with cells undergoing EMT. The effect of agrin on enhancing stress fibers and pFAK a focal adhesions as well as down-regulation of β-catenin at the cell-cell junction indicates agrin promotes EMT and an evident mesenchymal fate.

To confirm the in vitro effects of agrin on EMT and cell migration we used an immortalized mouse epicardial cell line MEC1 (Li et al., 2011) in a wound-healing assay. Addition of agrin, following an automated scratch assay, at 200ng/ml promoted wound healing at an equivalent rate to 5ng/ml TGFβ, with 75% of the scratch closed after 64 hours. Blocking antibody combinations against integrin α4β1 also promoted wound healing to the same level as agrin-treatment. In contrast treatment with an agrin blocking antibody decreased wound healing to 42%, compared with 55% following control PBS treatment. Interestingly, blocking antibody against the agrin binding partner dystroglycan severely reduced the mouse EPDC migration to 20%, and this effect could not be rescued by addition of TGFβ (**Figure S6**). These data suggests that agrin has comparable, but independent, effects to TGFβ in promoting EMT and migration of mouse epicardial cells resulting in enhanced wound closure.

### Agrin regulates EMT through aggregation of Dystroglycan to the Golgi apparatus maintaining basement membrane and cytoskeletal connectivity

To further assess a role for dystroglycan in agrin-mediated EMT, we investigated the distribution of dystroglycan in human epicardium-like cells treated following exogenous agrin treatment (**Figures 7A-7G**). In cells treated with DMSO dystroglycan was detected at low levels around nuclei (**Figure 7A**), whereas stimulation with 200ng/ml agrin led to aggregation of dystroglycan in dense perinuclear puncta (**Figure 7B**, white arrows). In contrast, blocking endogenous agrin with a neutralizing antibody resulted in the dissociation of dystroglycan puncta (**Figure 7C**, white arrow). Blocking dystroglycan with a DAG1 antibody had a similar effect to agrin antibody (**Figure 7D**, white arrow), suggesting agrin signals through dystroglycan in this cell model. Combined treatment of cells with agrin together with a DAG1 antibody or agrin antibody, partially rescued the dissociation of dystroglycan puncta from the nuclei (**Figures 7E, 7F**). In contrast with agrin, TGFβ did not aggregate dystroglycan around the nuclei despite promoting EMT of the human epicardial-like cells (**Figure 7G**).

**Figure 7.**
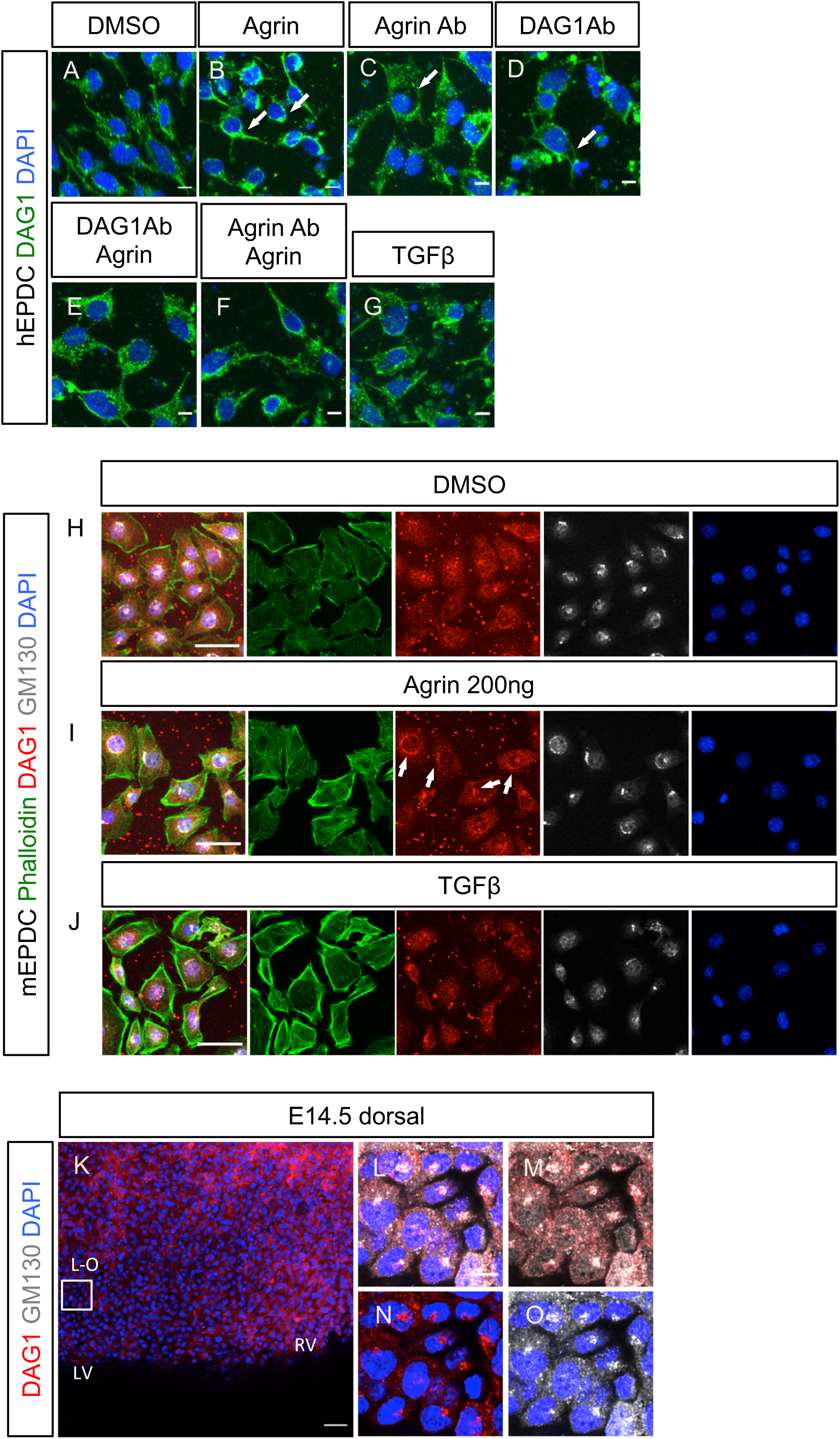
Agrin aggregates DAG1 to the Golgi apparatus in epicardial cells. (A-G) Immunofluorescent staining for DAG1 (green) and DAPI (blue) of human epicardial-like cells treated with agrin or blocking antibodies. Compared with the negative DMSO control (A), DAG1 was detected in proximity of nuclei when 200ng/ml agrin was added (B, white arrows). Blocking agrin with agrin blocking antibody resulted in dispersal of dystroglycan (white arrow in C). Treatment with blocking antibody against DAG1 also resulted in dystroglycan dispersal (white arrows in D). These effects were partially rescued by addition of agrin (E and F). TGFβ treatment did not aggregate dystroglycan proximal nuclei (G). (I-J): Immunofluorescent staining with DAG1 antibody and GM130 antibody showing agrin aggregates dystroglyan to the Golgi apparatus in epicardial cells. Addition of agrin and TGFβ both enhanced stress fiber shown with Phalloidin staining (green). Addition of agrin caused aggregation of dystroglycan (red, arrows in I) which overlapped with the Golgi apparatus labeled with GM130 (white). TGFβ did not aggregate dystroglycan (J). (K-O): Wholemount immunofluorescent staining in E14.5 hearts showing dystroglycan in epicardial cells aggregated in Golgi apparatus labeled with GM130. (K): dorsal view of E14.5 heart of Dystroglycan (red) and DAPI (blue). (L-O): magnified view of the inset box in (K). Scale bar: A-J:10 μm. K: 50 μm. L: 10 μm. LV: left ventricle. RV: right ventricle.

The subcellular pattern of Dystroglycan aggregation induced by agrin suggested localization to a discrete cell organelle(s). A previous report revealed that Golgi-resident glycosyltransferases fukutin and fukutin-related protein are required for post-translational modification of Dystroglycan (Esapa et al., 2002), suggesting dystroglycan more specifically may localize in the Golgi apparatus. To test this hypothesis, we investigated the localization of dystroglyan, following treatment with exogenous agrin, with the Golgi marker GM130 (Nakamura et al., 1995, 1997). Under basal/control conditions, dystroglycans were detected on the plasma membrane, with mild enrichment in the nuclear envelope in some cells (**Figures 7H**). Treatment with 200ng/ml agrin led to enhanced stress fibers (compare **Figure 7H and 7I**) and condensed aggregation of dystroglycans proximal to the nuclei. The localization of dystroglycan detected with DAG1 antibody fully overlapped with the GM130+ Golgi apparatus (**Figure 7I**). TGFβ treatment also enhanced the incidence of stress fibers (compare **Figure 7J and 7H**) but did not induce aggregation of dystroglycans on the Golgi apparatus (**Figure 7J**).

We further examined the morphology of Golgi apparatus in vivo within epicardial cells. In E14.5 hearts, GM130 labeled the Golgi apparatus as dense puncta within the developing epicardium. Dystroglyan, detected with DAG1 antibody, overlapped with GM130^+^ puncta indicating dystroglycan is also aggregated in the Golgi apparatus *in vivo* (Figure **7K-O**).

These data collectively identify dystroglycan as a key downstream component for agrin signaling in promoting epicardial EMT and suggest that dystroglycan enrichment in the Golgi apparatus is a key component of the mechanism of action of agrin in maintaining ECM basement membrane and epicardial cytoskeletal integrity.

## Discussion

In this study, we characterized the role of ECM components during heart development and specifically in the essential process of epicardial EMT. Regions of active EMT were associated with the deposition of specific matrix components in the epicardial layer. Specifically, we identified a novel and essential role for the basement membrane proteoglycan agrin during epicardial development. Our data revealed that agrin promotes epicardial EMT in both mouse and human models, through aggregation of dystroglycans to the Golgi apparatus, connecting extracellular signals from ECM with intracellular pathways required for epicardial cell activation. The function of ECM in development has been studied previously, but there is limited insight into ECM signaling and the regulation of cellular processes in the forming heart.

### Epicardial EMT and ECM

Using SEM, we observed morphological heterogeneity within epicardial cells across different development stages. In the E13.5 and E14.5 epicardium, we observed large flat cells tightly connected with their neighbors in addition to much smaller, round cells which were clearly separated from each other. The morphologies of these cells are consistent with the definition of epithelial versus mesenchymal cells (Yang et al., 2020), suggesting we have observed *de novo* epicardial EMT at high sub-cellular resolution. Recently, epicardial cells with similar morphologies were described and attributed to stages of maturity (Velecela et al., 2019); in contrast, our data provides novel insight into the co-existence of distinct cellular morphologies within the developing epicardium as correlated with EMT status.

Consistent with the observations from our SEM analyses, active and inactive regions of EMT were associated with the presence of key ECM components. We found that integrin α4 was prominently down-regulated in regions of high EMT; an observation supported by a previous report that knock-down of integrin α4 in chicken epicardial cells resulted in a highly invasive phenotype and increased migration of cells into the underlying myocardium of chicken-quail chimera (Dettman et al., 2006). However, loss of agrin in this study resulted in down-regulation of integrin α4 and decreased EMT, indicating that integrin α4 has additional roles beyond restraining epicardial cells. It is plausible that integrin α4 also acts as an anchoring molecule for epicardial cells to attach to their extracellular environment and serves to relay signals for proliferation and maintenance. Laminin, a major component of the basement membrane (reviewed by Yurchenco, 2015), was continuous and localized between the epicardial cells and the myocardium in inactive EMT regions. In contrast within active EMT regions, laminin was discontinuous and reduced among epicardial cell clusters suggesting loss of epicardium-myocardium connectivity via the basement membrane. The difference in the deposition of laminin and α4 integrin in the high-versus low-EMT regions suggests that the basement membrane and interstitial ECM are highly dynamic structures during epicardial development.

### Agrin is a novel regulator of epicardium development and EMT

The impact of agrin loss on the proliferation and migration of WT1^+^ cells indicates agrin is an essential regulator of epicardial development. Our SEM analysis of the agrin KO epicardial cells revealed several morphological changes compared to wild type littermates; including fewer protrusions at cell-cell junctions and on the cell surface. These protrusions have not been described before within the forming epicardium and although the have been described for other cell types their function remains unknown (Velecela et al., 2019, Figure 4). Based on our observation that cells undergoing EMT have more protrusions than cells with an epithelial morphology, and agrin KO epicardial cells have fewer protrusions than controls, we hypothesize that these protrusions are correlated with EMT, possibly as result of cytoskeletal remodelling. Loss of agrin compromised several ECM components within the epicardium, such as integrin α4, podoplanin. Laminin and collagen I were also down-regulated in agrin KO mutants, not only in the epicardium but also in the underlying myocardium, indicating a more global role for agrin in ECM organization. The up-regulation of cytosolic β-catenin in the epicardium and cells adjacent to the epicardium suggests decreased EMT potential in the mutant cells. Interestingly, in a previous study agrin was reported to stabilize the blood-brain barrier (BBB) by stabilizing the localization of tight junction proteins such as VE-Cadherin, β-catenin and ZO-1 at cell-cell junctions (Steiner et al., 2014). In contrast, in C2C12 myoblast, overexpression of β-catenin blocked the effect of agrin on the aggregation of the acetylcholine receptor (AchR) (Wang et al., 2008), suggesting an antagonistic relationship between agrin and β-catenin.

We observed coronary vasculature defects within the agrin KO hearts, which phenocopied those observed in WT1 mutants (Gise et al., 2011), as a possible consequence of compromised epicardial EMT. However, agrin is also an important component of the basement membrane of the vasculature. The abnormal arrangement of PECAM^+^ endothelial cells of the coronary vessel in agrin KO hearts is, therefore, more likely a direct consequence of agrin loss in blood vessel endothelium. Indeed, agrin can stabilize Vascular Endothelial Growth Factor Receptor-2 (VEGFR-2) in the ECM and thus promote angiogenesis (Njah et al., 2019).

### Agrin promotes EMT through aggregation of Dystroglycan

Agrin has multiple isoforms, including membrane-anchored and secreted variants (Burgess et al., 2000; Neumann et al., 2001). Agrin regulates intracellular signalling pathways through binding to the low-density lipoprotein receptor 4 (LRP4) and muscle-specific tyrosine kinase (MuSK), which has been characterized in detail in muscle-neuron junctions (Zhang et al., 2008). Dystroglycan is another known binding partner of agrin (Gee et al., 1994; Scotton et al., 2006) and is an essential component of the dystrophin-associated protein complex. Both α- and β-dystroglycans are encoded by the *DAG1* gene (Ibraghimov-Beskrovnaya et al., 1992). Dystroglycan links the extracellular matrix with the actin cytoskeleton (Ervasti and Campbell, 1993). In cardiomyocytes, dystroglycans associate with phosphorylated YAP, the Hippo pathway effector, and inhibit cardiomyocyte proliferation (Morikawa et al., 2017). Agrin promotes cardiomyocyte proliferation in post-myocardial infarction hearts through DAG1 and YAP (Bassat et al., 2017). Although dystroglycans are usually detected in the cell membrane, a nuclear localization signaling (NLS) sequence has been identified in β-Dystroglycan (Lara-Chacón et al., 2010). There is also report that dystroglycans can be translocated to the nuclear membrane in C2C12 myoblast cell lines, playing an important role in maintaining the integrity of the nuclear envelope (Gracida-Jiménez et al., 2017; Velez Aguirela et al., 2018). Our results suggest that agrin regulates epicardial EMT through aggregation of dystroglycans. Disaggregation and dispersal of dystroglycans was observed in agrin KO hearts coincident with impaired EMT, suggesting that the agrin-dystroglycan signaling axis is important for maintaining ECM/basement membrane and epicardial cell integrity and cytoskeletal cross-talk to ensure mesenchymal transition. Agrin may regulate downstream signaling pathways such as ERK through aggregated dystroglycans on the plasma membrane. In cultured epicardial cells as well as embryonic hearts, this aggregation appears to be via the Golgi apparatus. A recent study reported that Golgi compaction is related to the mesenchymal status of lung adenocarcinoma cells (Tan et al., 2017), implicating the dynamics of the Golgi apparatus as an important factor influencing EMT. Aggregation of dystroglycans is tightly correlated with the expression level of WT1 in epicardial cells further supporting a role in EMT. Dystroglycans must be glycosylated in the Golgi apparatus to acquire function (Michele et al., 2002), thus another possibility is that agrin regulates downstream signals through post-translational modification of dystroglycans.

In summary, our results identify agrin and its receptor dystroglyacan as important regulators of EMT in the embryonic epicardium and provide novel insights into the essential function of the ECM in the developing mammalian heart.

## MATERIAL AND METHODS

Key resources table

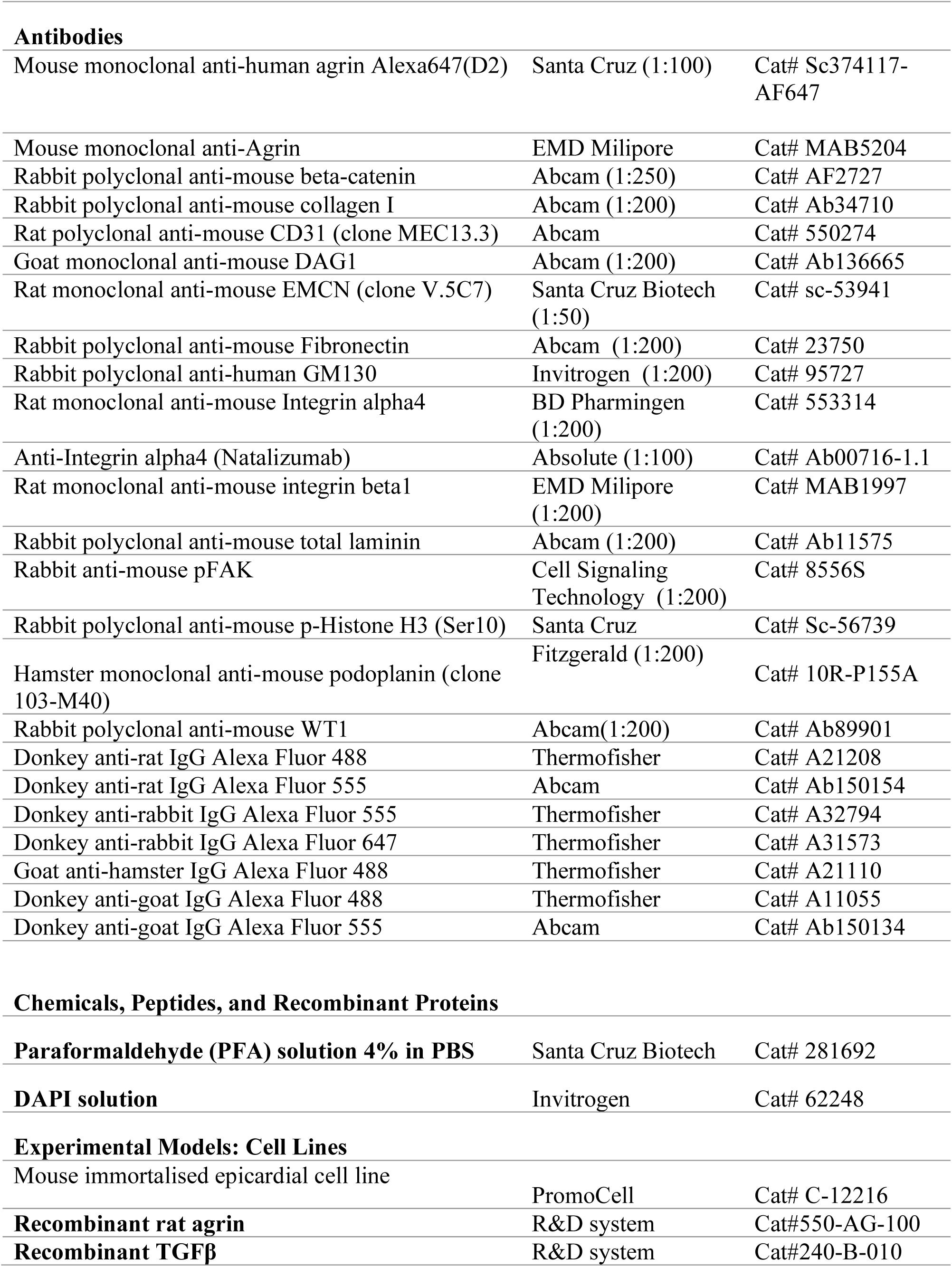
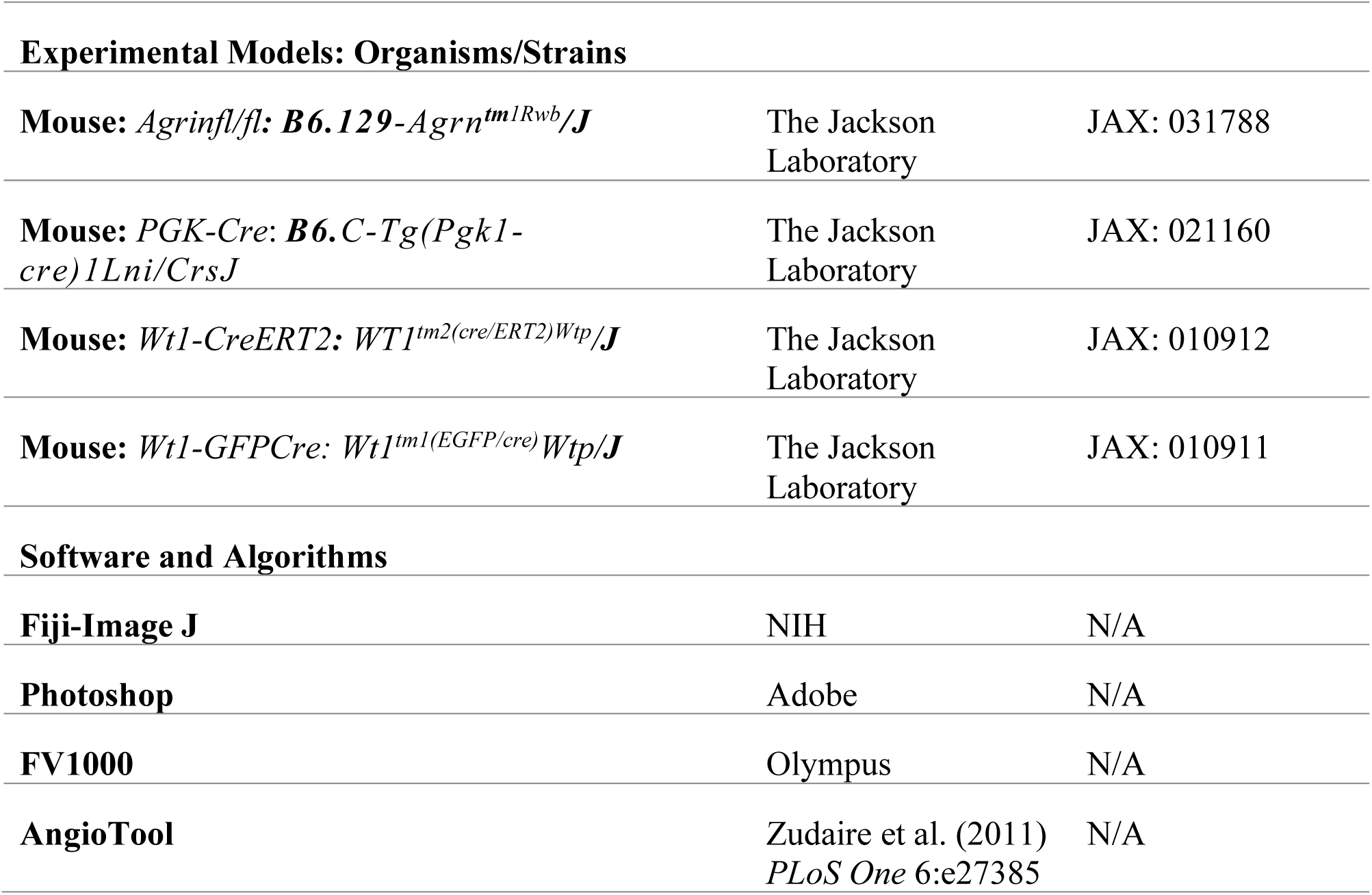

### Mouse lines

Genetically modified mouse lines were kept in a pure C57BL/6 background. *Agrinfl/fl* mice were crossed with *PGK-Cre* mice to generate *Agrin*^*+/-*^ progenies. *Agrin*^*-/-*^ (*AgrinKO*) mice were generated form intercrossing of *Agrin*^*+/-*^ males and females. *Wt1-CreERT2* mce were crossed with *Wt1-GFPCre* to generate *Wt1*^*KO*^ embryos. For timed-mating experiments, mice of breeding ages were paired in breeding cages and vaginal plugs were checked in the morning until the females were plugged. The date of a vaginal plug was set as embryonic day (E)0.5. Mice were housed and maintained in a controlled environment by the University of Oxford Biomedical Services. All animal experiments were carried out according to the UK Home Office project license (PPL) 30/3155 and PPDE89C84 compliant with the UK animals (Scientific Procedures) Act 1986 and approved by the local Biological Services Ethical Review Process.

### Scanning Electron Microscopy

Samples were fixed in 2.5%glutaraldhyde pH7.4 at 37°C after dissection then stored at 4°C overnight. After rinsed with PBS, the samples were postfixed with 1%OsO4 for 1 hr then rinsed with distilled water. After dehydration through ethanol gradient, the samples were dried with a critical point dryer. They were then mounted on a stub with carbon tape and coated with gold-sputter for imaging with the Zeiss Sigma 300 FEG-SEM operating at 2kV.

### Fluorescent Immunohistochemistry

Embryos were harvested at the required stages and their hearts were dissected in PBS and fixed with 4%PFA overnight. For wholemount staining, samples were washed in 0.3% TritonX-100 in PBS and blocked in 1%BSA 0.3% TritonX-100 in PBS for more than 2 hours. The samples were then incubated with the primary antibodies with the dilution as listed in the key source table in the blocking solution overnight at 4°C. In the second day, the samples were washed for more than five times in 0.3% TritonX-100 in PBS. The samples were then incubated with the secondary antibodies and DAPI diluted in PBS overnight at 4°C. The samples were then washed with PBS for more than 5 times the next day and mounted in 50% glycerol in PBS for scanning with Olympus FV1000 confocal microscope.

To prepare tissue sections, the fixed embryonic hearts were embedded in OCT compound embedding medium. Serial sections were prepared with a cryostat microtome. Cells or heart explants were cultured in 10%FBS DMEM on coverslips coated with 0.1%gelatin. For sections and cell cultures, glass slides or coverslips were washed twice with PBS to remove the OCT then permealise with 0.5% TritonX-100 in PBS for 10 min and washed with PBS afterwards. The sections or cells were then blocked in 10% donkey or goat serum, 1%BSA and 0.1%TritonX-100 in PBS for more than 1 hr at room temperature. Next, samples were incubated with primary antibodies diluted in the blocking solution at the concentration indicated above overnight at 4°C. The next day, samples were washed 3 times in 0.1% TritonX-100 in PBS for 10 minutes then incubated with corresponding fluorescent secondary antibodies diluted at 1:200 in 0.1% TritonX-100 in PBS for 1hour. After three washes with 0.1%TritonX-100 and 10min of DAPI staining, slides were mounted in 50% glycerol in PBS and imaged with Olympus FV1000 confocal microscope. Images were analysed with FIJI.

### hESC-derived Epicardium-like cell differentiation

Human embryonic stem cells (hESCs; H9 line) were cultured in mTeSR™1 basal medium (Stem Cell Technologies), in 6-well plates coated with matrigel (Corning Matrigel Matrix hESC-qualified). Once hESCs reached 75% confluency, the differentiation was initiated, based on the method described in the Iyer D. et al. study^1^, with certain modifications.

More specifically, for the CDM-PVA (Chemically Define Medium -PolyVinyl Alcohol) basal cardiac differentiation medium, human insulin was purchased by SIGMA and was used at a final concentration of 7ug/ml, while WNT3A (R&D Systems) was used at a concentration of 50ng/ml. Furthermore, on day 5 of differentiation, when the Lateral Plate Mesoderm cells were passaged to generate epicardium, the cells were seeded at a 75×10^4^/cm^2^ density with additional supplementation of 10uM Rock-inhibitor (Y-27632, Abcam). The next day Rock-inhibitor was removed by replenishing with the cardiac differentiation medium, CDM-PVA supplemented with WNT3A (50ng/ml), BMP4 (50ng/ml, R&D Systems) and all trans Retinoic Acid (4μM, SIGMA). The differentiation continued until day 14, with the media being replenished once more at day 11, and topped up with fresh solution as required when the colour of the cell media turned orange.

### Mouse EPDC EMT Induction

Embryonic immortalised mouse epicardium-derived cells (mEPDC) were treated with 2.0 µg/ml human recombinant transforming growth factor (TGF)-β (R&D Systems; 240-B) for 30 hrs to induce EMT. Cells were then fixed in 4% PFA prior IF assessment.

### Wound Heal Capacity

mEPDC cells wound heal capacity was assessed by initiating a scratch wound in confluent cell monolayer using the 96-pin WoundMaker™ (Essen BioScience), and wells were washed 2x with media to remove floating cells. Immediately following wounding, cell treatments were added in growth medium and the plate incubated at 37 °C, 5 % CO_2_. Wound images were taken every 4 hrs, relative wound density was analysed using the live content imaging system IncuCyte HD (Essen BioScience).

### Statistical analysis

All data are presented as mean ± standard error of the mean (SEM). Statistical analysis was performed on GraphPad Prism 8 software. The statistical significance between two groups was determined using an unpaired two-tailed Student’s *t*-test, these included an F-test to confirm the two groups had equal variances. Among three or more groups, one-way analysis of variance (ANOVA) followed up by Tukey’s multiple comparison test was used for comparisons. A value of p ≤ 0.05 was considered statistically significant.

### Contact for reagent and resource sharing

Further information and requests for resources and reagents should be directed to and will be fulfilled by the Lead Contact, Paul R. Riley.

## Acknowledgements

This work was supported by the British Israel Research and Academic Exchange (BIRAX 13BX14PRET) and a Leducq Foundation Transatlantic Network of Excellence Program 14CVD04 to XS and the British Heart Foundation (chair award CH/11/1/28798 and programme grant RG/08/003/25264 to PRR) and supported by the BHF Oxbridge Centre of Regenerative Medicine (RM/13/3/30159). KK is supported by a Wellcome Trust Four year PhD Studentship 215103/Z/18/Z to KK; JMV is supported by a BHF Intermediate Basic Science Research Fellowship FS/19/31/34158. We thank Errin Johnson of Sir William Dunn School, University of Oxford for support on SEM; Alain Wainman, Andrew Jefferson and Nadia Halidi of Micron, University of Oxford for support on confocal. We thank staff at the Oxford Biomedical Service Building for animal husbandry.

