## Supplementary Figures for "The extracellular matrix protein agrin is essential for epicardial epithelial-to-mesenchymal transition during heart development"

### Supplementary Figure 1

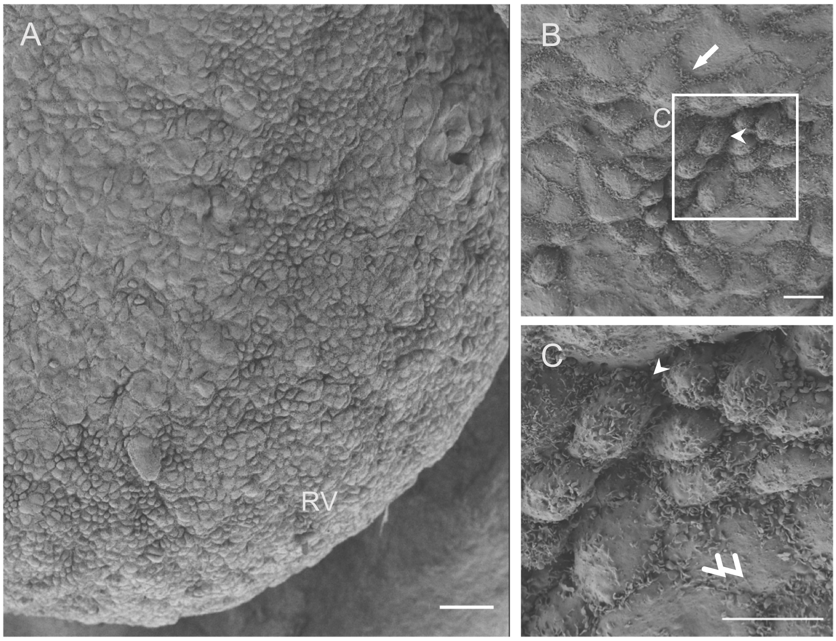

**Supplementary figure 1. Cells with epithelial morphology and mesenchymal morphology within the embryonic epicardium.** (A): the dorsal surface of an E13.5 embryonic heart, showing epicardial cells with different morphologies. (B): Magnified epicardium focusing on large and flat, epithelial-like cells (white arrow) and small, pillar-shaped, dispersed mesenchymal-like cells (white arrowhead). (C): magnified view of the inset box from (B). Scale bar: A, 50 μm. B, C: 10 μm. RV: right ventricle.

Supplementary Figure 2

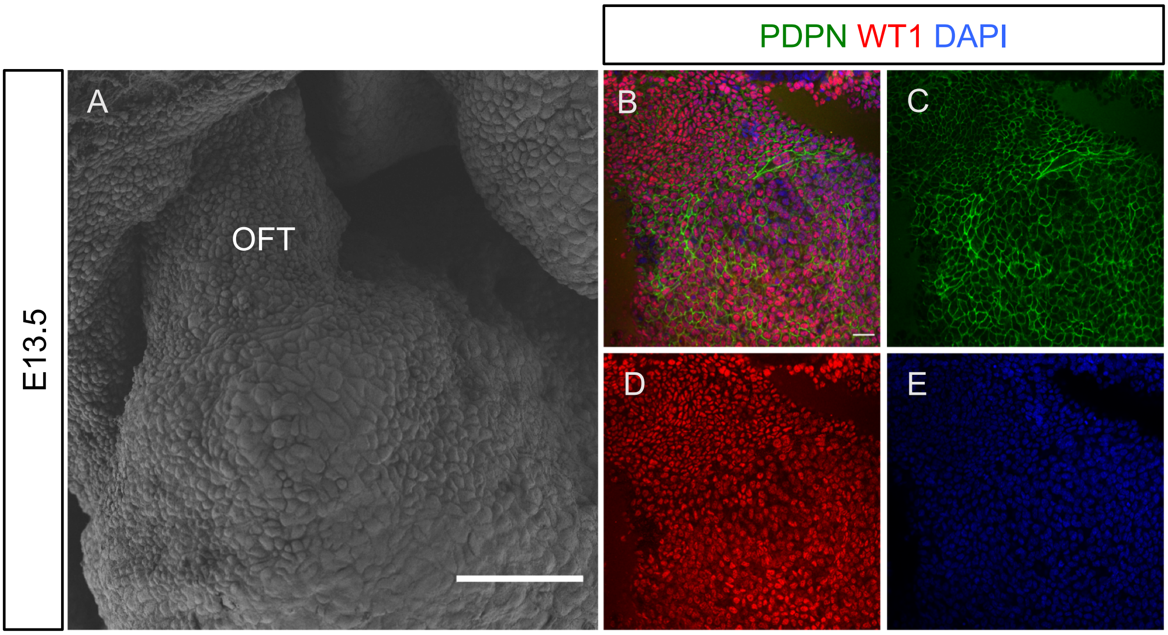

**Supplementary figure 2. Morphological differences between epicardial cells in the OFT.** (A): the OFT of an E13.5 heart under SEM: note the presence of distinct small, round cells and large, flat cells. (B-E), Whole-mount immunofluorescent staining of E13.5 for podoplanin (PDPN; green) and WT1 (red). Scale bar: 50 μm. OFT: outflow tract.

### Supplementary Figure 3

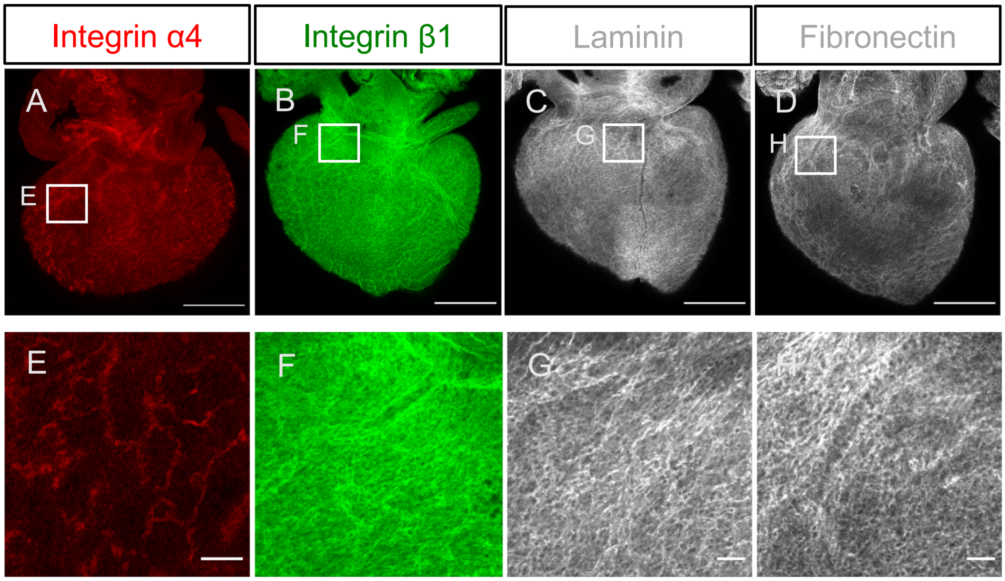

**Supplementary figure 3. Expression of extracellular matrix components within the embryonic epicardium.** (A-D). Wholemount immunofluorescent staining of integrin α4 (red), integrin β1 (green,), laminin (white) and fibronectin (white) on E14.5 embryonic hearts. E-H: magnified view of the inset boxes from A-D. Scale bar: A-D, 500 μm. E-H: 50 μm.

### Supplementary Figure 4

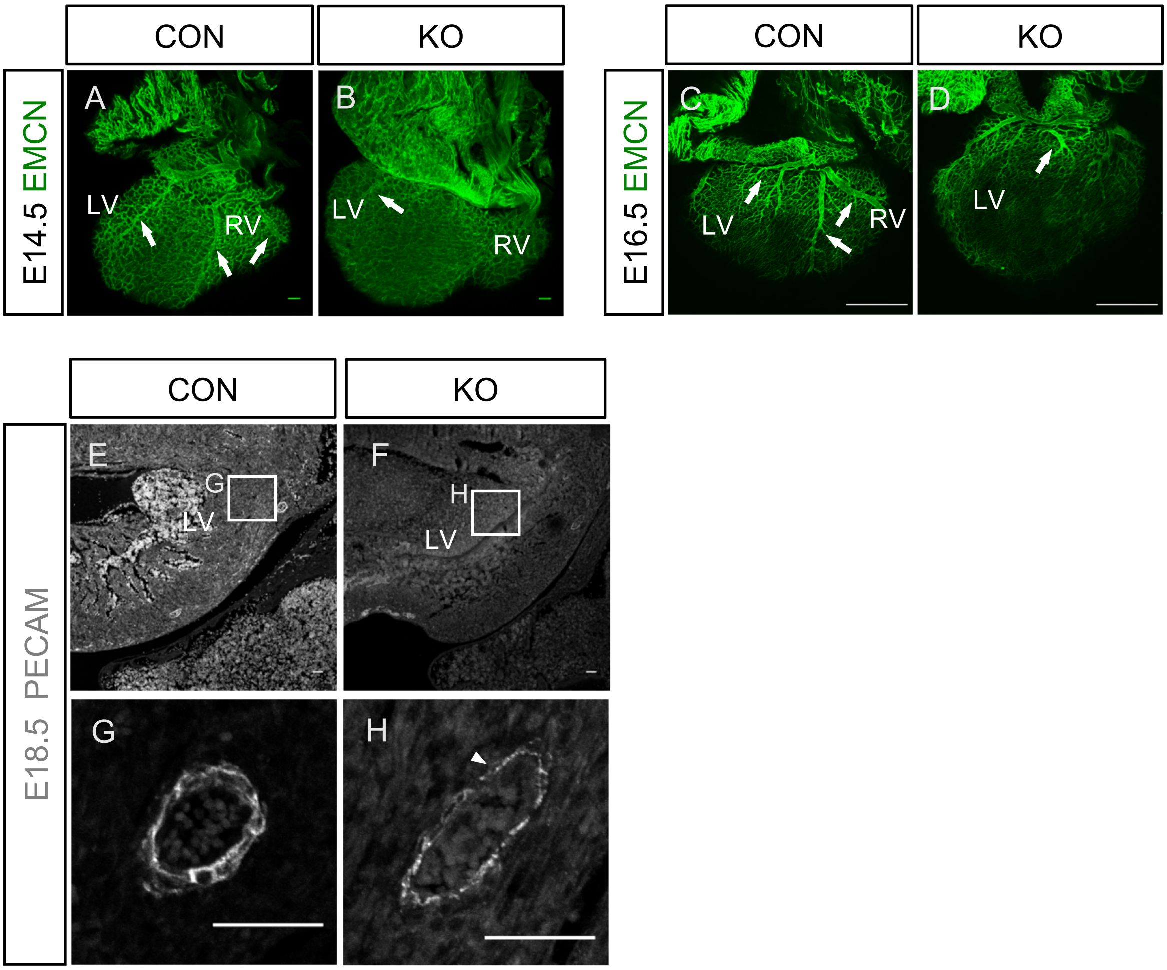

**Supplementary figure 4. Coronary vasculature defects in agrin KO hearts.** (A-D), whole-mount immunofluorescent staining for EMCN (green) of E14.5 (A, B) and E16.5 (C, D) littermate controls (A, C) and agrin KO hearts (B, D) labeling the coronary vasculature. Dorsal aspects are presented. Note the clearly defined and extended major vessels in the control heart (white arrows in A) which are virtually absent in the KO heart (white arrow in B). In E16.5 agrin KO heart, the main vessels are truncated (white arrows in D). (E-H), immunofluorescent staining on sections from E18.5 litter control hearts (E, G) and agrin KO (F, H) hearts for PECAM labeling endothelial cells. (G, H) are magnified views from the inset boxes in (E, F) showing a transverse section of a main coronary vessel. Note the weak and discontinuous PECAM staining in the agrin KO heart indicative of vessel instability (white arrow in H). Scale bar: A, B: 100 µm, C, D: 500 µm. E-H: 50µm.

### Supplementary Figure 5

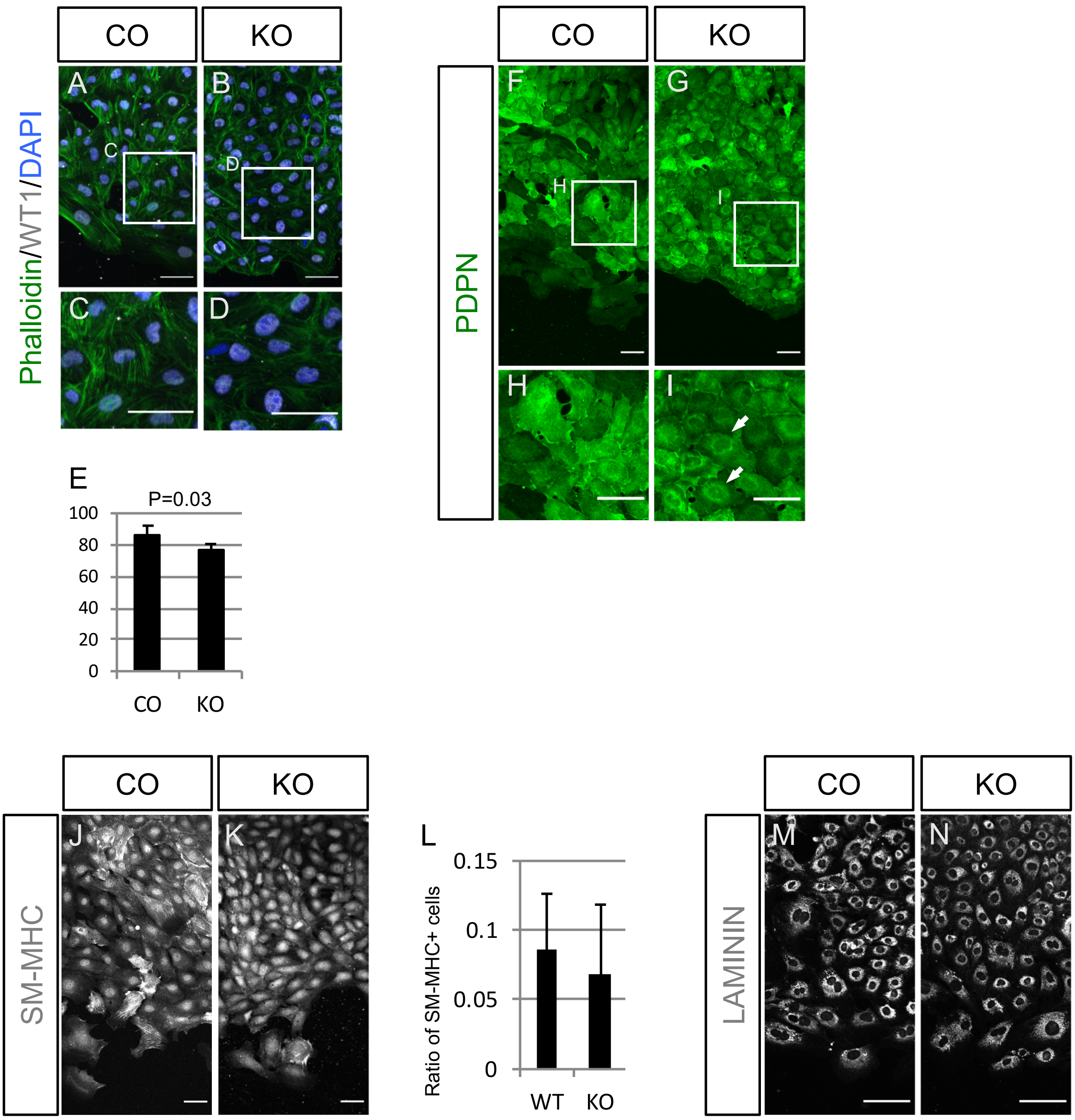

**Supplementary figure 5. Migration and ECM defects in epicardium-derived cells from agrin-KO explants.**

(A-B) Immunofluorescent staining of epicardium-derived cells derived from E11.5 explants of littermate control and agrin KO for cell stress fibers (phalloidin, green), WT1 (white) and nuclei (DAPI). (C, D) are magnified view of the inset boxes in (A, B). (E) Quantification of $WT1^{+}$ cells in control explant-derived and agrin KO derived epicardial cells. Data represent mean ± SEM. N=4 hearts per group. Significant differences (p value) were calculated using an unpaired, two-tailed Student's *t*-test. (F, G) Immunofluorescent staining for podoplanin (green) in epicardium-derived cells from control and agrin KO explants. (H and I) are magnified view of the box in D and E. Epicardial cells derived from the agrin KO explant showed weak podoplanin on the cell surface, especially in peri-nuclear regions (white arrows in I). (J, K, M, N) Immunofluorescent staining for SM-MHC (J, K) and Laminin (M, N) of epicardial cells from littermate control (J, M) and agrin KO explants (K, N). (L): quantification of SM-MHC-positive cells in total explant-derived epicardial cells. Data represent mean ± SEM; n=4 independent samples per group. Significant differences (p value) were calculated using an unpaired, two-tailed Student's *t*-test.

All scale bars: 50 µm.

Supplementary Figure 6

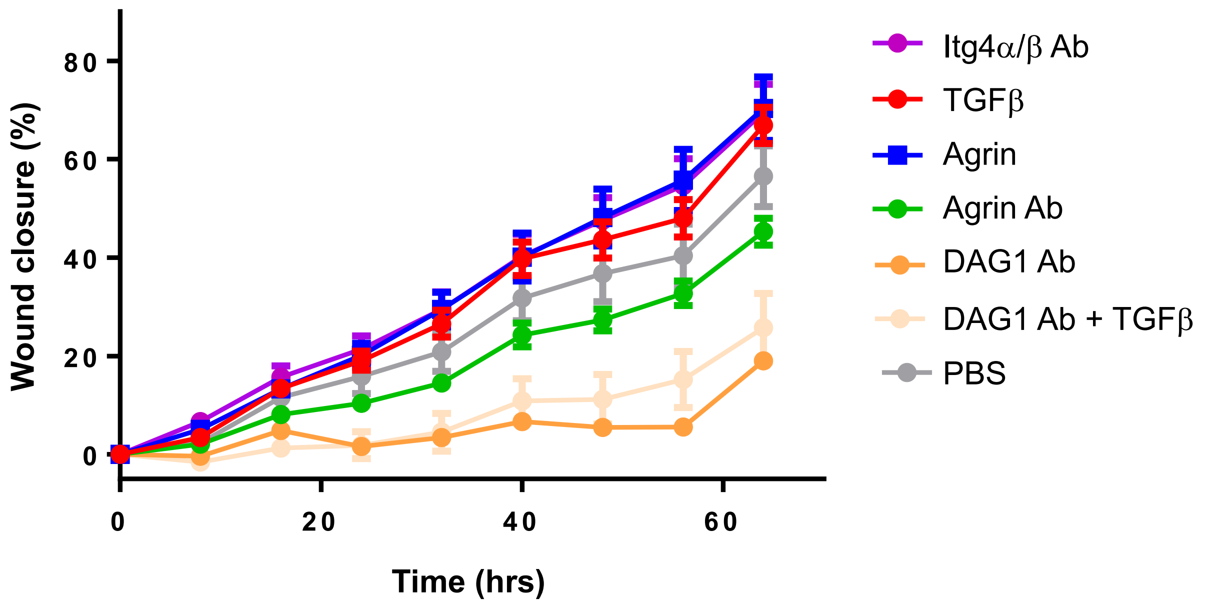

**Supplementary figure 6. Agrin enhances the migration of mouse epicardial cells.** Immortalised mouse epicardial cells were cultured for 64 hours with the indicated reagents and the wound closure was evaluated. Data represent mean ± SEM; n=12 independent treatments per group. Significant differences (p value) were calculated using an unpaired, two-tailed Student's *t*-test. Agrin treatment significantly increased the rate of 75%wound closure compared to untreated (PBS) and even TGFβ-treatment.
